# Optimizing the use of gene expression data to predict plant metabolic pathway memberships

**DOI:** 10.1101/2020.07.15.204222

**Authors:** Peipei Wang, Bethany M. Moore, Sahra Uygun, Melissa D. Lehti-Shiu, Cornelius S. Barry, Shin-Han Shiu

## Abstract

Plant metabolites produced via diverse pathways are important for plant survival, human nutrition and medicine. However, the pathway memberships of most plant enzyme genes are unknown. While co-expression is useful for assigning genes to pathways, expression correlation may exist only under specific spatiotemporal and conditional contexts. Utilizing >600 expression values and similarity data combinations from tomato, three strategies for predicting membership in 85 pathways were explored: naive prediction (identifying pathways with the most similarly expressed genes), unsupervised and supervised learning. Optimal predictions for different pathways require distinct data combinations that, in some cases, are indicative of biological processes relevant to pathway functions. Naive prediction produced higher error rates compared with machine learning methods. In 52 pathways, unsupervised learning performed better than a supervised approach, which may be due to the limited availability of training data. Furthermore, using gene-to-pathway expression similarities led to prediction models that outperformed those based simply on gene expression levels. Our study highlights the need to extensively explore expression-based features and prediction strategies to maximize the accuracy of metabolic pathway membership assignment. We anticipate that the prediction framework outlined here can be applied to other species and also be used to improve plant pathway annotation.

## Introduction

Metabolites are products of pathways consisting of a linked series of chemical reactions mainly catalyzed by enzymes (Berg et al., 2002). Plants produce diverse metabolites essential for plant growth, development, survival and adaptation (Stitt et al., 2010). Some of these metabolites are also important for human nutrition and medicine (Schmidt et al., 2007; Martin and Li, 2017). Gaining knowledge of metabolic pathways is important for understanding the rules of life and harnessing the metabolic potential of organisms through breeding and engineering. Although there is an increasing body of knowledge related to plant metabolic pathways (Verpoorte, 1998; Pichersky and Gang, 2000; Kim and Buell, 2015), many genes responsible for the biosynthesis of plant metabolites in known pathways remain to be identified and there are likely unknown pathways yet to be discovered (De Luca et al., 2012; Schlapfer et al., 2017). A key challenge in understanding plant metabolic pathways is that genes encoding metabolic enzymes have undergone large scale duplications in plant genomes and tend to exist as members of large gene families, such as the BAHD acyltransferase (Yu et al., 2009), cytochrome P450 (Yu et al., 2017), 2-oxoglutarate dependent dioxygenase (Kawai et al., 2014), and UDP-glycosyltransferase (Wilson and Tian, 2019) families. In addition, experimental assessments of metabolic pathway membership using biochemical and genetic approaches can be laborious. Therefore, it is important to prioritize candidate genes for functional analyses using computational predictions.

Substantial effort has been devoted to predicting metabolic pathway membership of plant enzyme genes. One approach is to first identify candidate enzyme genes from a plant genome, and then assign these enzyme genes into pathways according to the potential reactions the enzymes may participate in (Chae et al., 2014; Schlapfer et al., 2017). This approach was utilized to develop the Plant Metabolic Network database, a community resource, which is currently populated with pathway annotations from 125 plant and green alga species. Here the prediction of candidate enzyme genes is mainly based on sequence similarity of unknown genes to experimentally evaluated enzyme genes in model species.

Another approach to predict metabolic pathway genes is to associate unknown genes with existing pathway genes by exploring gene expression similarity based on the assumption that genes from the same pathway, have similar expression profiles (Segal et al., 2003; Kim and Buell, 2015). Considering the ease of generating transcriptome data in model and non-model species, gene expression data is an important resource for computational inference of gene function. There are three general strategies that are used to leverage expression data for functional inference. The first strategy, referred to as “naive prediction,” is to ask, for a gene of unknown function, which genes with known functions have the highest expression similarities to that gene. Despite its simplicity, the utility of this strategy has been shown in a number of single-gene studies (Hirai et al., 2007; Peng et al., 2015; Righetti et al., 2015; Huang et al., 2019), but its accuracy on a genome-wide scale is not clear. The second strategy is to globally infer functional association between plant genes through unsupervised machine learning approaches, where genes are first grouped into co-expression clusters and then genes of unknown function are assigned functions based on the identities of genes with known functions that are over-represented within clusters. This strategy is by far the most widely used (Mutwil et al., 2011; Serin et al., 2016; Uygun et al., 2016; Wisecaver et al., 2017; Abu-Jamous and Kelly, 2018; Gupta and Pereira, 2019). The third strategy is supervised machine learning approaches where the function of a gene is predicted with models learned from the expression profiles of genes with known functions. While supervised learning has been applied to predict functions in other contexts (Lan et al., 2007; Kaundal et al., 2010; Libbrecht and Noble, 2015; Lloyd et al., 2015; Ni et al., 2016; Moore et al., 2019), it has not been applied to predict plant metabolic pathway membership using gene expression data. It is currently unknown which of these two machine learning strategies is more effective and whether their accuracy varies depending on the pathway.

In addition to uncertainty about specific computational strategies, it is also unresolved how gene expression data should be used in pathway membership prediction. For example, it has been shown that, rather than using as many different conditions as possible (Aoki et al., 2016; Obayashi et al., 2018), expression similarities measured using distinct subsets of expression data can be more accurate for inferring functional relationships for specific pathways (Usadel et al., 2009; Uygun et al., 2016; Wisecaver et al., 2017). Thus, in addition to choosing a computational strategy, it is important to optimize the use of specific expression data in metabolic pathway prediction. To accomplish this, we utilized tomato as a model organism due to the considerable knowledge of the diverse metabolic pathways in this species and the availability of a large collection of gene expression data (Carrari and Fernie, 2006; Mathieu et al., 2009; Brasher et al., 2015; Matsuba et al., 2015; Fan et al., 2017; Leong et al., 2019). We investigated the effect of expression datasets, expression values (absolute expression values or fold change), and gene expression similarity measures on the ability to predict pathway membership using three strategies: naive prediction, unsupervised learning, and supervised learning.

## Results & Discussion

### Relationship between gene expression similarity and metabolic pathway membership

We first examined the extent to which genes within the same metabolic pathway have similar expression profiles. Toward this end, we collected annotation data of 2171 tomato genes from 297 pathways with ≥ 5 annotated genes each (**Supplemental Data Set 1,2**), and transcriptome data from 372 experiments (each experiment includes multiple biological/technical replicates, **Supplemental Data Set 3,4**) (see **Methods**). Using Pearson’s correlation coefficient (PCC) calculated using enzyme gene expression levels (Fragments Per Kilobase Million, FPKM) from all experiments as the expression similarity measure, we found that for 124 out of 297 pathways (41.8%) there was significantly higher gene expression similarity between pairs of genes from the same pathway than between pairs of genes randomly chosen from different pathways (Wilcoxon signed-rank test, *p* < 0.05; **Figure 1A**, **Supplemental Data Set 5**). When the maximum expression similarity between a gene and all other genes in the same pathway was examined, genes in 202 of 297 pathways (68.0%) had significantly higher expression similarities within a pathway than between pathways (**Supplemental Figure 1A**, **Supplemental Data Set 5,6**). This finding was not simply due to the contribution of paralogs that might have higher expression similarities than non-paralogous pairs (**Supplemental Figure 1D**). These results suggest that expression data can be useful for distinguishing genes within pathways from genes between pathways. In addition, the maximum expression similarity within a pathway was more useful for distinguishing genes within and between pathways than the median similarity (**Supplemental Figure 1B,C**). However, even using maximum PCC as a criterion, 72.7% of genes did not have above-threshold PCC values (PCC=0.66, the 95^th^ percentile value of between-pathway gene pairs) within a pathway (**Supplemental Figure 1A**). Therefore, we asked how similarity measures should be generated to best identify pathway membership.

**Figure 1.**
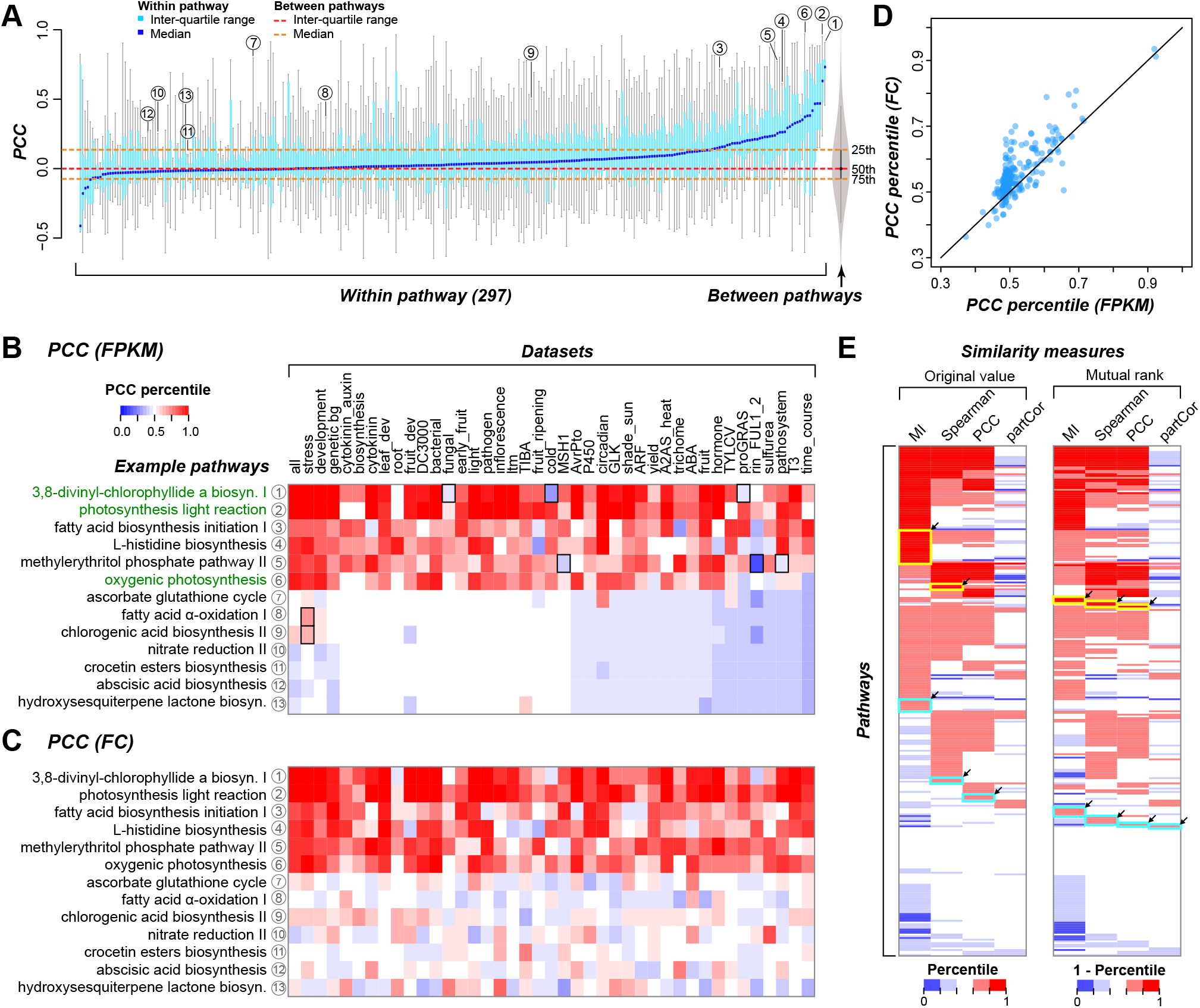
Impact of expression dataset, expression values, and similarity measures on expression similarities between genes in the same pathway. (**A**) Gene expression similarity (in terms of PCC) within and between pathways. Boxplot showing expression similarity between genes within individual pathways. Blue line: median value; light blue box: interquartile range. Gray violin plot shows the PCC distribution of all gene pairs between different pathways (between-pathway distribution), where median value and interquartile range are marked with red and orange dashed lines, respectively. Circled numbers indicate example pathways in (**B**). (**B**) Examples showing the effect of dataset on gene expression similarity. FPKM was used to calculate the PCCs. X-axis: 41 datasets, y-axis: example pathways. Color scale: percentile of the median PCC within a pathway in the between-pathway distribution (percentile_BP_). The percentile_BP_ values were scaled to 0–1 here and hereafter, where 1 indicates the 100^th^ percentile. Pathway names in green are those relevant to photosynthesis. (**C**) Same as (**B**), except that FC was used in PCC calculation. (**D**) Scatter plot showing the differences between percentile_BP_ of median PCC calculated using FPKM (x-axis) and FC (y-axis). (**E**) Gene expression similarity calculated using different similarity measures. Color scale: percentile_BP_ of median expression similarity calculated using different similarity measures (left) and 1 □ percentile_BP_ of the mutual rank of similarity values (right). For mutual rank, 1 □ percentile_BP_ was used because lower ranks (lower percentiles) indicate higher degrees of expression correlation. Yellow and cyan rectangles indicate pathways with high (red) and median high (light red) expression similarities, respectively, only when a specific similarity measure was used.

For any gene pair, expression similarity is influenced by three factors: (1) which expression dataset is used, e.g., all data or a subset; (2) what expression value is used, i.e., expression levels or contrasts in the form of fold change; and (3) which similarity measure is used, e.g., PCC vs. Spearman’s Rank. For a pair of genes in a pathway, we were not interested in their absolute expression similarity. Rather, we were interested in how their similarity was different from a distribution of between-pathway gene similarities that we treated as the background, null distribution. Thus, in all subsequent analyses, we determined the percentile value of each within-pathway similarity in the between-pathway similarity distribution (hereafter referred to as percentile_BP_). The median of percentile_BP_ values for all within-pathway pairs was calculated, and a higher median percentile_BP_ value indicates that genes in that pathway had higher similarity to each other compared with between-pathway pairs. In the above analysis, all 372 experiments were used. To evaluate whether different datasets may be more useful for determining expression similarities of genes in different pathways, 41 datasets including 36 from individual studies, and five combined sets consisting of samples from (1) tissue/developmental stage/circadian clock experiments, (2) different genetic backgrounds (e.g., mutants), (3) different hormone treatments, (4) different stress treatments, and (5) all 372 experiments (**Supplemental Data Set 3,4**, see **Methods**) were examined.

### Example pathways demonstrating the effect of expression datasets used on expression correlations among pathway genes

Using FPKM as the expression value and PCC as the similarity measure, the datasets that produced the highest percentile_BP_ for the 13 example pathways differed substantially (**Figure 1B**, results for all the pathways in **Supplemental Data Set 7**). Some pathways, such as those relevant to photosynthesis (green, **Figure 1B**, **Supplemental Data Set 7**), tended to have high PCC percentile_BP_ values for most datasets used. For example, expression of genes in the 3,8-divinyl-chlorophyllide a biosynthesis I pathway (labeled ⍰ in **Figure 1B**) was well correlated in all datasets except fungal inoculation, cold treatment and treatment with paclobutrazol and gibberellic acid (proGRAS dataset, **Supplemental Data Set 3**), consistent with the finding that photosynthesis is disrupted under multiple stress conditions (Nouri et al., 2015). Another example is the methylerythritol phosphate (MEP) pathway II (⍰ in **Figure 1B**) where pathway genes were generally well correlated in expression except when the *FRUITFULL1/2* (*FUL1/2*) and *RIPENING INHIBITOR* (*RIN*) mutant dataset, MutS HOMOLOG1 (MSH1) gene silencing dataset and the Pseudomonas inoculation dataset were used. Note that genes in the carotenoid biosynthesis pathway downstream from MEP are differently regulated by FUL1/2 and RIN (Fujisawa et al., 2014). This indicates that genes in MEP II may also be differently regulated by FUL/1/2 and RIN; when these transcriptional regulators are mutated or silenced, MEP II genes may not be properly regulated and thus not co-expressed. In contrast to the first six examples shown in **Figure 1B**, genes in the remaining seven example pathways are either correlated in a highly dataset-specific manner or not correlated in any dataset. For example, genes in the fatty acid α-oxidation I pathway (⍰, **Figure 1B**) and chlorogenic acid biosynthesis II pathway (⍰, **Figure 1B**) only had appreciable levels of expression correlation when the combined dataset with different stress treatments was used, consistent with the role of these two pathways in protecting plants against environmental perturbations such as oxidative stress (De Leon et al., 2002; Niggeweg et al., 2004). These results demonstrate the need to consider datasets that best reflect the biological processes in which different pathways participate.

Using fold change (FC) values from the same datasets (see **Methods**), instead of absolute expression levels (FPKMs), led to improved percentile_BP_ values for all example pathways in the corresponding between-pathway distribution (each of the 41 datasets had a dataset-specific between-pathway PCC distribution, calculated using FC) (**Figure 1C**, **Supplemental Data Set 8**). In fact, 250 of 297 pathways had a higher percentile_BP_ when using FC than when FPKM was used (**Figure 1D**), demonstrating the influence of the format of the expression values used for analysis. We should note that the crocetin ester biosynthesis pathway in crocus (Carmona et al., 2006) and the 3β-hydroxysesquiterpene lactone biosynthesis pathway in feverfew (Majdi et al., 2011) (⍰ and ⍰ in **Figure 1B,C**) are not expected to be present in tomato. Our finding of beyond the randomly expected degree of co-expression between genes in these pathways when FC values were used (**Figure 1C**) likely reflects the existence of similar pathways in tomato, which need to be further verified experimentally.

### Impact of expression similarity measure on expression correlations among pathway genes

Thus far, we used PCC as a similarity measure to assess linear correlation. Because gene expression correlation can be non-linear or context dependent, we explored three additional similarity measures: Spearman’s rank, Mutual information (MI), and partial correlation (partCor). MI measures the mutual dependence between two variables (Steuer et al., 2002), while partCor measures the degree of association between two variables with the effect of other factors removed (Wang, 2013). The percentile_BP_ values based on these three new measures were calculated as previously performed for PCC. Because the inference of gene function association when using similarity measures, such as PCC values, can be affected by the method of gene expression database construction and the target gene function (Obayashi and Kinoshita, 2009), we also determined the mutual rank (MR, calculated as the geometric mean of the rank of expression similarity between gene1 and gene2 in all expression similarities between gene1 and all other genes and the rank in all expression similarities between gene2 and all other genes, see **Methods**) for each of the four measures. For MR, 1 minus the percentile_BP_ was calculated because gene pairs with higher percentile_BP_ will have smaller numerical rank values.

To illustrate the impact of different similarity measures on expression correlations among pathway genes, we used the FPKM values of genes in all experiments and found that genes in some pathways only displayed high similarity values when specific measures were used (**Figure 1E**, **Supplemental Data Set 9**). For example, the expression similarity scores of genes from 19 pathways were in the highest quintile (red) only when MI was used (left panel, **Figure 1E**). Using MR of MI, an additional four pathways now had gene expression similarities in the highest quintile (right panel, **Figure 1E**). Genes in pathways with higher quintiles for all four measures generally tended to have higher similarities in both expression level and breadth (**Supplemental Figure 2**). On the other hand, genes in pathways with higher PCC/Spearman quintiles but lower MI/partCor ones had similar patterns in expression levels but not in expression breadths (**Supplemental Figure 2**). In contrast, genes in pathways with higher MI quintiles compared with other measures had higher correlation in expression breadth than expression level (**Supplemental Figure 2**). Taken together, these findings highlight the importance of exploring expression datasets, expression values, and similarity measures in evaluating gene expression similarity for different pathways. Thus, in subsequent analyses, we determined expression correlations using 41 datasets, 2 types of expression values, and 8 similarity measures to yield a total of 656 possible data combinations.

### Naive prediction of pathway genes

With the effects of expression dataset, expression value (i.e., FPKM vs. FC), and similarity measure demonstrated, we next asked what approaches should be used to predict whether a gene belongs to a metabolic pathway based on expression similarity. We explored three general approaches: (1) naive prediction, (2) unsupervised learning, and (3) supervised learning. The latter two approaches are detailed in subsequent sections. The first approach was named naive prediction solely because it is the simplest approach. We explored two methods for assigning a gene to a pathway: 1) naive median: If gene X has the highest median expression similarity with genes in pathway A, then gene X is predicted to be in pathway A; and 2) naive maximum: If gene X has the maximum expression similarity with gene Y and gene Y is in pathway B, then gene X is predicted to be in pathway B (**Figure 2A**). Genes annotated to multiple pathways were excluded from analysis because they would be only assigned to a single pathway using naive prediction approaches; after excluding these genes, 972 enzyme genes were assigned to 85 pathways, each with ≥ 5 genes (**Supplemental Data Set 10**).

**Figure 2.**
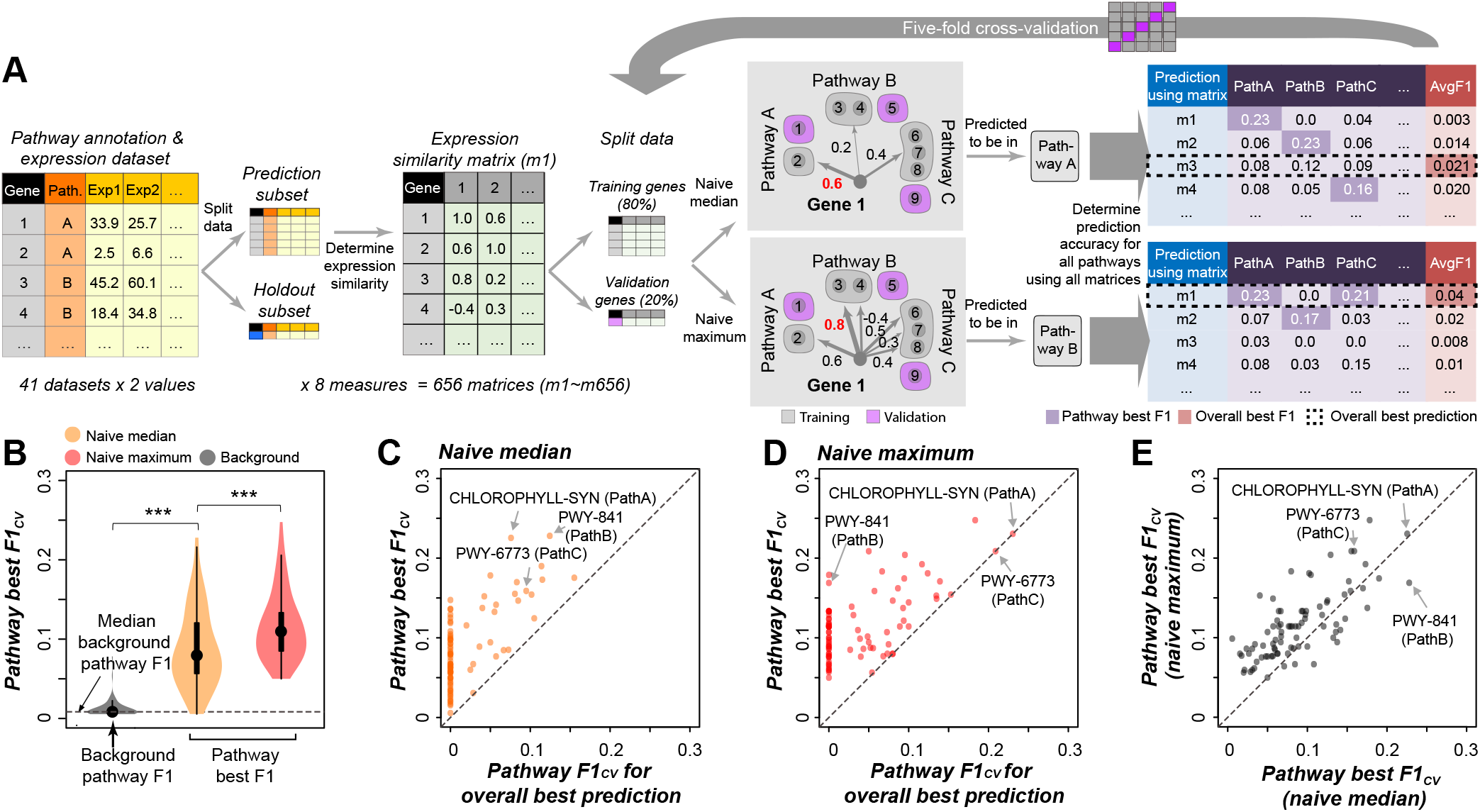
Naive prediction of metabolic pathway genes. (**A**) Methodology. For each of the 82 (41 expression datasets × 2 expression values) expression matrices, five genes (blue) were randomly held out from each pathway containing ≥ 25 genes. The remaining data (gray) were used to determine expression similarities among genes using 8 similarity measures, resulting in 656 expression similarity matrices (m1–m656). Genes within each pathway were further split into “training” (80%, gray) and “validation” (20%, magenta) subsets, and the data splitting was conducted five times. The example validation gene 1 is predicted to be in pathway A using the naive median method because it has the highest median expression similarity with training genes in pathway A; however, it is predicted to be in pathway B using the naive maximum method because it has the maximum expression similarity with gene 3, which belongs to pathway B. The thickness of the arrow and number beside the arrow indicate the degree of expression similarity. All 656 expression similarity matrices were used for both methods, and the F1_CV_ score was calculated for each of the 85 pathways, resulting in two 656 x 85 F1 score matrices. The prediction with the highest F1_CV_ for a pathway (purple) was referred to as the pathway best prediction. The average F1_CV_ across 85 pathways for each naive model(made using one of the 656 matrices) was calculated to measure the overall prediction performance, and the model with the highest average F1 (red) was referred to as the overall best model (dashed box). (**B**) Distribution of pathway best F1_CV_ obtained using the naive median (orange) and naive maximum (pink) methods. Distribution of background pathway F1 scores is shown as a gray violin plot, with the median value indicated by a dashed line. ***: *p*-value < 0.001, Wilcoxon signed-rank test. (**C,D**) Performances of naive median (**C**) and naive maximum (**D**) models. X-axis: F1_CV_ values for the overall best model—values in the dashed box in the table on the right in (**A**). Y-axis: pathway best F1_CV_ across all 656 models—values in the purple cells in the table on the right in (**A**). (**E**) Comparison of pathway best F1_CV_ for naive median (x-axis) and naive maximum (y-axis) predictions. Three example pathways in (**C-E**) show the differences in pathway predictions made by the two approaches, and are shown in the table on the right in (**A**) as Pathway A (CHLOROPHYLL-SYN, 3,8-divinyl-chlorophyllide a biosynthesis I), B (PWY-841, superpathway of purine nucleotides de novo biosynthesis I), and C (PWY-6773, 1,3-β-D-glucan biosynthesis).

The goal for pathway prediction is ultimately to predict pathway membership of unknown genes. Thus, we split annotated genes in a pathway into “known” and “unknown” subsets. The “known” genes, referred to as “*training*” genes, were used for establishing prediction “models”, which were simply the naive median and naive maximum rules for naive predictions. In the meantime, the “unknown” genes, for which annotation information is available, were used to validate the models in a five-fold crossvalidation scheme (referred to as “*validation*” genes, see **Methods**). For six pathways with ≥ 25 genes, we also held out five genes, referred to as “*test*” genes, prior to training-validation split to further test the models. To make the naive predictions comparable with predictions made using unsupervised and supervised learning approaches discussed in later sections, the same training-validation splits were used. To evaluate model predictions, the F1-score — the harmonic mean of precision (the proportion of predictions that are true positives) and recall (the proportion of true cases that are correctly predicted) — was calculated for each pathway based on prediction of cross-validation (CV) genes (referred to as F1_CV_) in each of the five data splits. The average F1 among the five data splits was used to evaluate the prediction performance (**Figure 2A**). F1 ranges from 0 and 1, with 1 indicating a perfect model. For the naive approaches, the end results of the calculations were two F1_CV_ score matrices, one for the naive medium and one for the naive maximum, that were each 656 (number of data combinations) by 85 (number of pathways) in dimension (**Supplemental Data Set 11,12**).

We first asked whether one data combination was particularly useful for predicting pathway membership (overall best, red box, **Figure 2A**). The best data combination had an average F1_CV_ = 0.04 (all experiments, FPKM as the expression value, MR of PCCs as the similarity measure, using the naive maximum method). Although better than the median F1 of random guess (0.01, dotted line, **Figure 2B**, and **Supplemental Data Set 13**), the prediction based on the best data combination is far from perfect. Because different data combinations affect whether pathway memberships can be recovered using expression data (**Figure 1**), we next assessed the extent to which the best data combination differs between pathways. By identifying the data combination that led to the highest F1_CV_ scores for each pathway (pathway best F1_CV_, purple boxes, **Figure 2A**), we found that genes in a pathway were better predicted when the pathway-specific optimal data combination was used; the median pathway best F1_CV_ values using the naive median and the naive maximum methods were 0.08 and 0.11, respectively (**Figure 2B**); both values were higher than that for the overall best data combination (0.04). In nearly all cases, the pathway F1_CV_ values obtained using the overall best data combination were substantially lower than the pathway best F1_CV_ values for both the naive median (**Figure 2C**) and the naive maximum (**Figure 2D**) methods. One notable pattern is that the overall best data combination frequently led to pathways with F1_CV_=0 (**Figure 2C,D**), further demonstrating the need to identify the optimal combinations of datasets, expression values, and similarity measures.

We also found that the naive maximum method generally performed better than the naive median (**Figure 2E**), consistent with our finding that the percentile_BP_ for the maximum expression similarity of a gene to other genes in the same pathway was higher than that for the median expression similarity (**Supplemental Figure 1**). This is likely because genes in the same pathway can be regulated at many other levels beyond the transcriptional level, thus resulting in relatively lower median expression similarity within pathways. In addition, different components of a pathway may branch off to other pathways, further contributing to regulatory differences within each pathway (Ihmels et al., 2004). Nonetheless, even for the naive maximum method, the median value of pathway best F1_CV_ among the 85 pathways was only 0.11. Thus, despite the usefulness of the naive prediction approach for assigning enzyme genes to pathways, there remains substantial prediction errors (**Supplemental Figure 3**). Therefore, we explored whether pathway predictions could be improved using machine learning approaches.

### Prediction of pathway genes using unsupervised learning methods

Unsupervised learning in the form of clustering is one of the most widely used methods to aggregate genes with similar functions. While unsupervised learning methods have been used to predict metabolic pathway memberships (Wisecaver et al., 2017), earlier studies used one or a few datasets without exploring the effects of using different datasets and types of expression values. To explore the effect of clustering algorithms and parameters on prediction performance, we focused on two algorithms with “predict” functions, *k*-means and Affinity Propagation (see **Methods**). For each algorithm and each of the 82 data combinations (41 datasets and two expression values), two types of input matrices were generated for clustering. The first type was simply the expression value matrix (referred to as Set A, **Figure 3A**). The second type was generated by first determining the expression similarities of each gene to other genes and then calculating the median and maximum similarities of pathways to the gene in question (Set B, **Figure 3B**).

**Figure 3.**
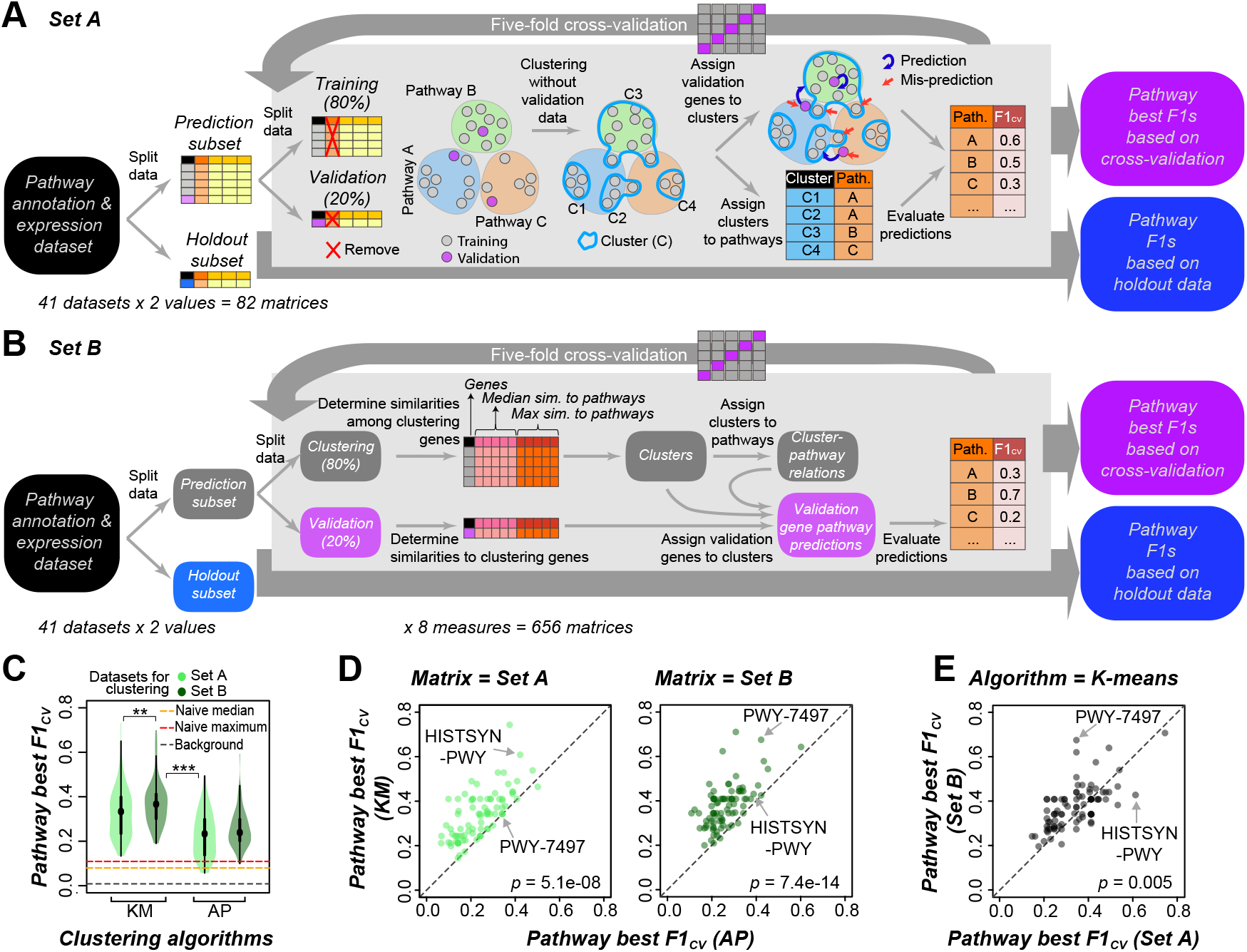
Prediction of pathway genes using unsupervised clustering methods. (**A**) Workflow for clustering using gene expression profiles in the form of 82 matrices (41 expression datasets x 2 expression values, Set A). Data splitting was conducted in the same way as for naive prediction approaches (*Split data* and *five-fold cross-validation* steps). The pathway annotations (colored circles, i.e., Pathways A, B, and C) were removed from the expression matrices before clustering. Clusters (blue closed lines, C1–C4) built using expression values of “training” genes (“*Clustering without validation data*”) were assigned to pathways based on enrichment analysis (“*Assign clusters to pathways*”, see **Methods**). Then the validation genes were assigned to clusters (see **Methods**), and then were assigned to the pathways the clusters were assigned to (“*Assign validation genes to clusters*”). Red arrow indicates mis-prediction of a gene. The average F1 scores of validation genes across the five training/validation splits (F1_CV_) were calculated to evaluate the predictions. The test genes were assigned to pathways using the same method used for the validation genes, and the average F1 of test genes from five clustering models (F1 based on holdout data, F1_test_) were calculated. (**B**) Similar to (**A**), except that clusters were built using gene-to-pathway co-expression matrices (Set B), i.e., the maximum and median expression similarity of a training gene to all other training genes in each pathway, which was calculated using eight different similarity measures, resulting in 656 (82×8) matrices and clustering models. For a validation or test gene, the expression similarity was calculated between the gene and the training genes in each pathway, and the maximum and median values were used in the matrix. (**C**) Distribution of pathway best F1_CV_ obtained from clustering models, performed using the *k*-means (KM, light green) or Affinity Propagation (AP, green) method, and the expression matrix (Set A) or gene-pathway co-expression matrix (Set B). Orange, pink and gray dashed lines indicate the median pathway best F1_CV_ from naive median and naive maximum prediction models, and the median background pathway F1 (as in **Figure 2B**), respectively. **: *p*-value < 0.01, ***: *p*-value < 0.001, Wilcoxon signed-rank test. (**D**) Comparison of pathway best F1_CV_ for *k*-means and Affinity Propagation clustering models, using Set A (left panel) or Set B (right panel). Dots: individual pathways. Two examples showing performance differences between two clustering methods: HISTSYN-PWY (L-histidine biosynthesis) and PWY-7497 (3β-hydroxysesquiterpene lactone biosynthesis). *p*-values are from Wilcoxon signed-rank test. (**E**) Comparison of pathway best F1_CV_ for *k*-means models using Set A or Set B data.

There are two potential issues complicating the prediction of pathway membership using clustering results. First, genes from multiple pathways can be grouped in the same cluster. Second, genes from a single pathway can be classified into several clusters. These issues were addressed by, for each cluster C, determining the enrichment scores for genes in every pathway and tentatively assigned C to a pathway, P, if the enrichment score of P for C was the highest among pathways and the enrichment was significant (see **Methods**). To alleviate the issue of overfitting, a fivefold cross-validation scheme was applied (**Figure 3A,B**, see **Methods**). This approach also allowed the clustering-based predictions to be compared with those based on supervised learning, which is detailed in the next section. To make predictions, each gene in the validation set was assigned to a cluster C if the distance between the gene in question and the cluster C centroid was the shortest among the distances between the gene and other cluster centroids, and then the gene was predicted to be in the pathway cluster C was assigned to. Because these clusters were used in making predictions, they are referred to as “clustering models”.

By comparing the F1_CV_ (the average F1 value for prediction on validation subsets across the five-fold cross-validation) from unsupervised learning with the F1_CV_ from naive approaches, we found that, the clustering-based unsupervised learning outperformed naive approaches by a large margin (**Figure 3C**), no matter which algorithm (*k*-means or Affinity Propagation) or dataset (Set A—gene expression value matrix, or Set B—gene-to-pathway expression similarity matrix) was used. This can be because the naive approaches only considered one expression correlation value (maximum or median), while the unsupervised approach utilized multiple expression correlation values between a gene in question to many other genes in multiple pathways. Nevertheless, like the naive approaches, no single clustering model had good predictions for most of the pathways; the maximum average F1_CV_ was only 0.07 (**Supplemental Figure 4**), which was obtained when the FPKM values of genes in the dataset with different genetic backgrounds were used in *k*-means clustering models (referred to as *k*-means models) with *k*=500. We also found that *k*-means outperformed Affinity Propagation for most pathways (**Figure 3D**), either using Set A (67 pathways) or Set B (77 pathways). In addition, *k*-means models using Set B outperformed that using Set A for 57 of 85 (67%) pathways (**Figure 3E**), which could be attributed to two potential reasons. First, there were too few features in some Set A datasets for clustering; the median number of features (expression values or contrasts) was 8 among 82 Set A datasets, whereas each Set B dataset had 170 features (median and maximum gene-to-pathway expression similarities among 85 pathways). Consistent with this hypothesis, there was a significant positive correlation (Spearman’s rho = 0.47, *p*-value = 6.4e-6) between the pathway best F1_CV_ and the number of features in the corresponding Set A dataset (**Supplemental Figure 5**). Second, gene-to-pathway expression similarity likely provided more information for pathway membership prediction than the gene expression profiles for most pathways. This may be because the gene-to-pathway expression similarity leverages structures of the metabolic pathway/network, which were captured by the unsupervised methods we used.

### Prediction of pathway genes using supervised learning methods

Different from unsupervised methods, where pathway information is not used for clustering, supervised machine learning methods build predictive models by learning from pathway annotations. In our first attempt to use supervised learning, the pathway prediction was framed as a multi-class learning problem where the goal was to predict which of the 85 pathways (i.e., classes) a gene belongs to using the Random Forest (RF) algorithm (see **Methods**). The same feature datasets (Set A and Set B) used for the unsupervised learning methods were used to build RF models (**Figure 4A,B**). Because supervised learning methods were used to directly associate the pathway labels with the underlying data, our expectation was that supervised learning models would outperform clustering-based predictions. Contrary to our expectation, the supervised learning models had an overall lower performance compared with clustering-based methods (**Figure 4C**, **Supplemental Data Set 13**). The median pathway best F1_CV_ values for RF models were 0.23 and 0.3, when Set A and Set B data were used, respectively, compared with 0.33 and 0.37 for *k*-means models.

**Figure 4.**
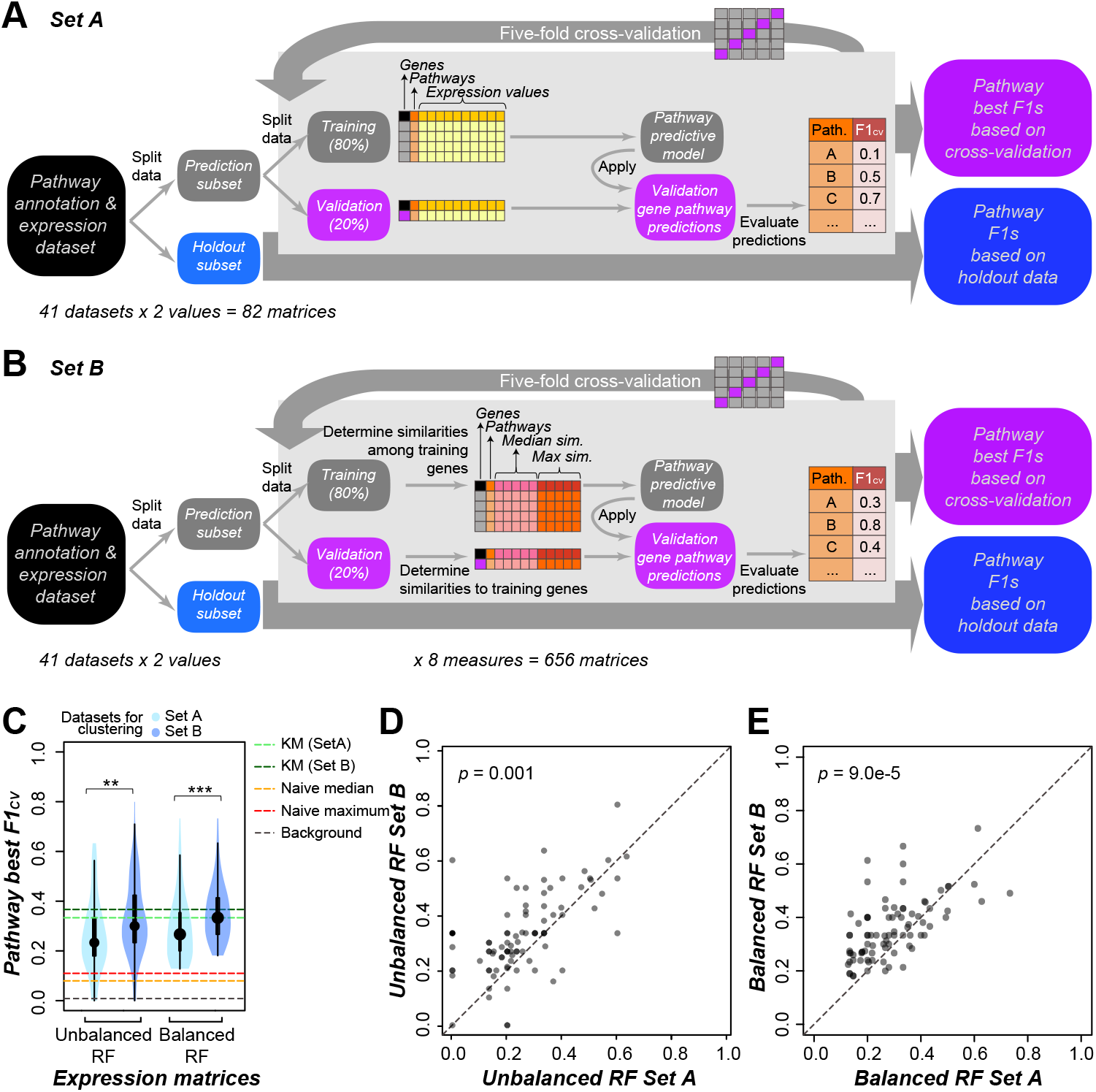
Prediction of pathway genes using supervised machine learning methods. (**A**) Workflow for RandomForest (RF) machine learning using Set A as features. Gene pathway membership information (i.e., labels) was merged with the expression matrix Data splitting was conducted in the same way as for naive and unsupervised approaches (*Split data* and *five-fold cross-validation*) in **Figure 2, 3**. A grid search was conducted to get the best combination of parameters (hyperparameters) that yielded the maximum F1_CV_ (see **Methods**). The final model was built using the same crossvalidation scheme with the hyperparameters and was applied to validation and test genes (“*Apply*”). (**B**) Same as (**A**), except that gene-to-pathway expression similarity matrices (Set B) were used as features. (**C**) Distribution of pathway best F1_CV_ from unbalanced RF or balanced RF models using Set A (light blue) or Set B (blue). Light green and green dashed line: median pathway best F1_CV_ from *k*-means models using Set A and Set B, respectively (as in **Figure 3C**); orange and pink dashed line: median pathway best F1_CV_ from naive median and naive maximum prediction models, respectively (as in **Figure 2B**); gray dashed line: median background pathway F1. **: *p*-value of Wilcoxon signed-rank test < 0.01. (**D, E**) Comparison of pathway best F1_CV_ from unbalanced RF (**D**) and balanced RF (**E**) models when Set A and Set B were used. Dots: individual pathways. *p*-value is from Wilcoxon signed-rank test.

One potential reason for the difference in performance is that there were too few genes in most pathways (median pathway size=8 even after filtering out small pathways, see **Methods**) for supervised machine learning to effectively generalize the features shared by genes in a pathway. Consistent with this, we found that the differences between pathway best F1_CV_ values of RF models and *k*-means models were weakly, but significantly correlated with pathway size, with Spearman’s rho = 0.41 (*p*-value=1.1e-4) and 0.28 (*p*-value=0.01) when Set A and Set B were used, respectively (**Supplemental Figure 5**). These results suggest that when there are too few genes in each pathway, clustering-based approaches may be superior to supervised learning methods. Nonetheless, the correlations between pathway size and F1_CV_ differences were weak, suggesting that other factors also explain the difference in performance. Upon examination of the confusion matrices (or error matrices, showing the proportion of genes in a pathway that are predicted as being in each of the 85 pathways) for *k*-means and RF models (**Figure 5A,B, Supplemental Data Set 14,15**), we found that a major reason for the poorer performance of RF models was that genes from most pathways tended to be mispredicted to be in two pathways: 260 genes from 64 pathways and 84 genes from 42 pathways were mispredicted to be in triacylglycerol degradation (LIPAS-PWY) and homogalacturonan degradation (PWY-1081), respectively. These mis-predictions could be attributed to the fact that LIPAS-PWY and PWY-1081 were the two largest pathways (75 and 54 genes, respectively, **Supplemental Data Set 10**) and jointly contributed to 12.6% of the training instances among 85 pathways. These unbalanced pathway data led to a bias in the model that ultimately contributed to the mis-predictions.

**Figure 5.**
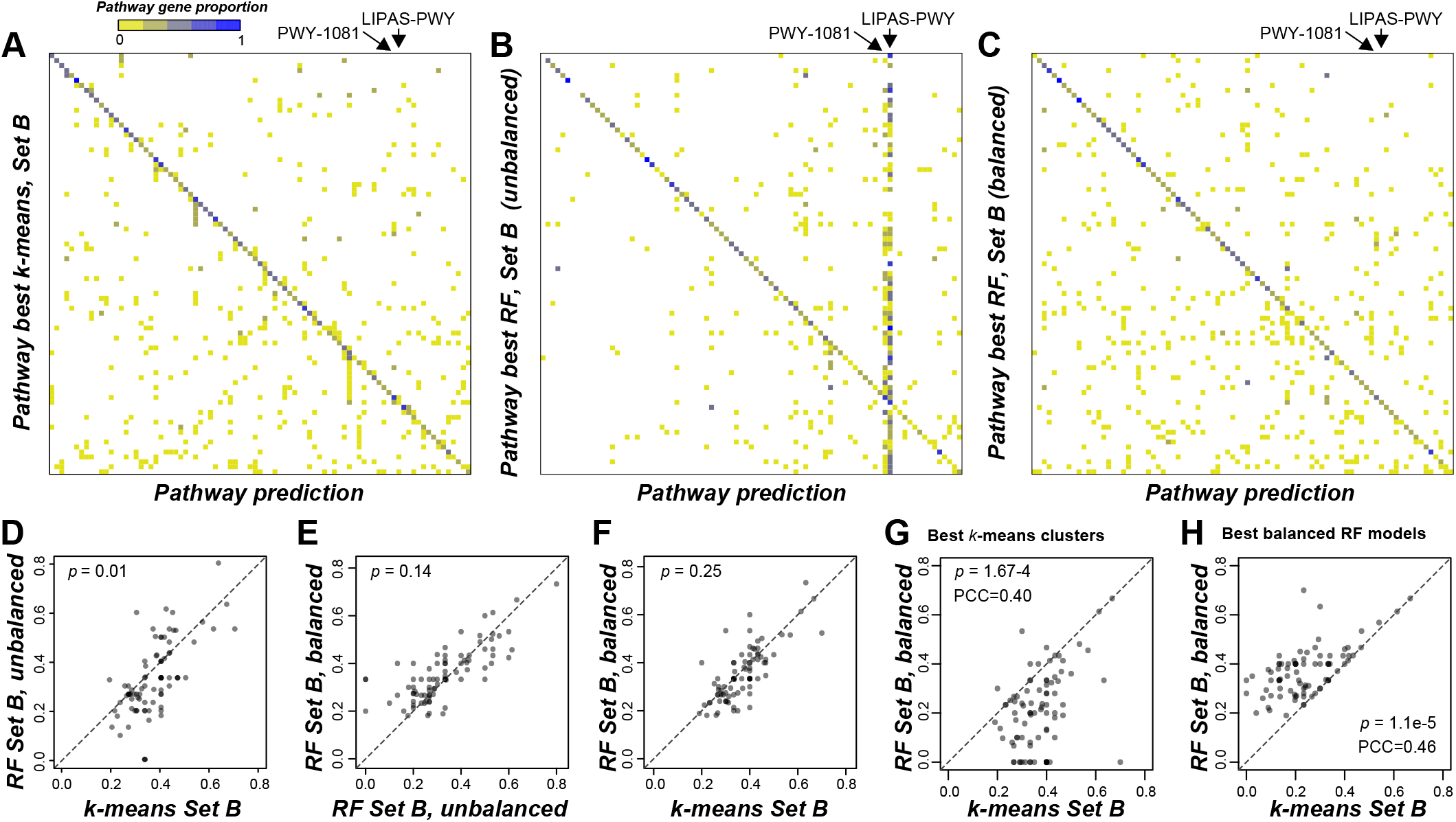
Performance difference between *k*-means clustering and Random Forest models when Set B was used. (**A-C**) Confusion matrix, which shows the proportion of genes that are predicted in each pathway for pathway best *k*-means models (**A**), unbalanced RF models (**B**) and balanced RF models (**C**). Color scale: proportion of genes in a pathway (y-axis) predicted as being in one of the 85 pathways (x-axis) by the pathway best model. (**D-F**) Comparison of pathway best F1_CV_ between *k*-means, unbalanced RF and balanced RF models when Set B was used. *p*-value is from Wilcoxon signed-rank test. (**G-H**) Pathway F1_CV_ from *k*-means (x-axis) and balanced RF models (y-axis) when the pathway best *k*-means Set B data (**G**) or the pathway best balanced RF Set B data (**H**) was used. Dots: individual pathways.

To alleviate the impact of sampling bias on model performance, we balanced the training data by randomly up-sampling the minority pathways to the size of the largest pathway in the training subset so that all 85 pathways had the same size (n = [75-5] x 0.8 = 56) in the training subset (see **Methods**). The instances in the validation subset were kept unchanged; thus the performance on the validation subset was still comparable with the performances of *k*-means and the original RF models based on unbalanced data. This approach led to new, balanced RF models. In contrast to unbalanced RF models, only six and 15 genes from three and nine pathways were mispredicted as in LIPAS-PWY and PWY-1081, respectively, and the prediction errors were relatively evenly distributed across pathways (**Figure 5C, Supplemental Data Set 16**). The resulting median pathway best F1_CV_ for balanced RF Set B models was 0.33, which was higher than that of unbalanced RF Set B models (0.30), while still slightly lower than that of *k*-means Set B models (0.37) (**Figure 4C**, and **Figure 5D-F**). Thus, only balanced RF models are further discussed.

Consistent with the observations from naive prediction and *k*-means models, the optimal combination of expression datasets, expression values and similarity measures used in RF models was different for each pathway (**Supplemental Figure 4**). Next, we asked whether the performances of *k*-means and RF models were similarly affected by data combinations, i.e., whether the best data combination for a pathway was the same for *k*-means and RF models. Because Set B data generally led to better predictions (**Figure 3E, Figure 4E**), only *k*-means and RF models using Set B (contains 656 datasets) were examined. Surprisingly, the best data combination for *k*-means and RF models was the same for only 16 of the 85 pathways (**Supplemental Data Set 17**). This result suggested that different information from the datasets were utilized in these two approaches. Although the best dataset differs for *k*-means and RF models, their prediction performances for the same data are significantly correlated (**Figure 5G,H**). Together with the observation that the pathway F1_CV_ values for all the *k*-means models were positively correlated with those for the RF models (PCC = 0.55, *p* < 2.2e-16, **Supplemental Figure 6**), these results suggest there are some commonalities in how *k*-means and balanced RF models learn from the gene-to-pathway expression similarity.

### Features important for pathway-specific predictions and likely causes of mispredictions in supervised learning methods

While we found that supervised learning models had a slight performance disadvantage compared with unsupervised clustering, supervised learning models allowed us to ask what expression features were the most important, i.e., contributed the most to predicting memberships in different pathways. This contribution was quantified using feature importance from the RF algorithm (see **Methods**). Here we focused on the balanced RF models using Set B data. Note that in Set B data, the predictive features for each gene were how well the expression of that gene was correlated to the expression of genes in each of the 85 pathways. The correlation was quantified as either the median or maximum gene-to-pathway expression similarity. Thus, there were 170 features ranked by their importance from 1 (best) to 170 (worst). We hypothesized that for a gene in pathway P, its expression similarity to other genes in pathway P (gene-to-target-pathway) should be more important for predicting that the gene belongs to pathway P than its similarity to genes in other, non-P pathways (gene-to-other-pathway). As expected, the highest ranking gene-to-target-pathway features for the pathway best RF models had significantly better (smaller) ranks than gene-to-other-pathway features (Wilcoxon signed-rank test, *p*-value = 1.8e-3, **Figure 6A**). In addition, the pathway best F1_CV_ was negatively correlated with the importance rank of the gene-to-target-pathway features (Spearman’s rho = −0.24, *p*-value = 0.02, **Figure 6B**). However, the correlation was not robust. In addition, in 81 out of 85 pathways, the most important features were in fact not the gene-to-target-pathway features. These findings indicated that even though gene-to-target-pathway similarities can be useful, relationships between genes and other pathways in terms of expression are also required. This also explains why naive prediction models based solely on gene-to-target-pathway expression correlation performed so poorly (**Figure 2B**).

**Figure 6.**
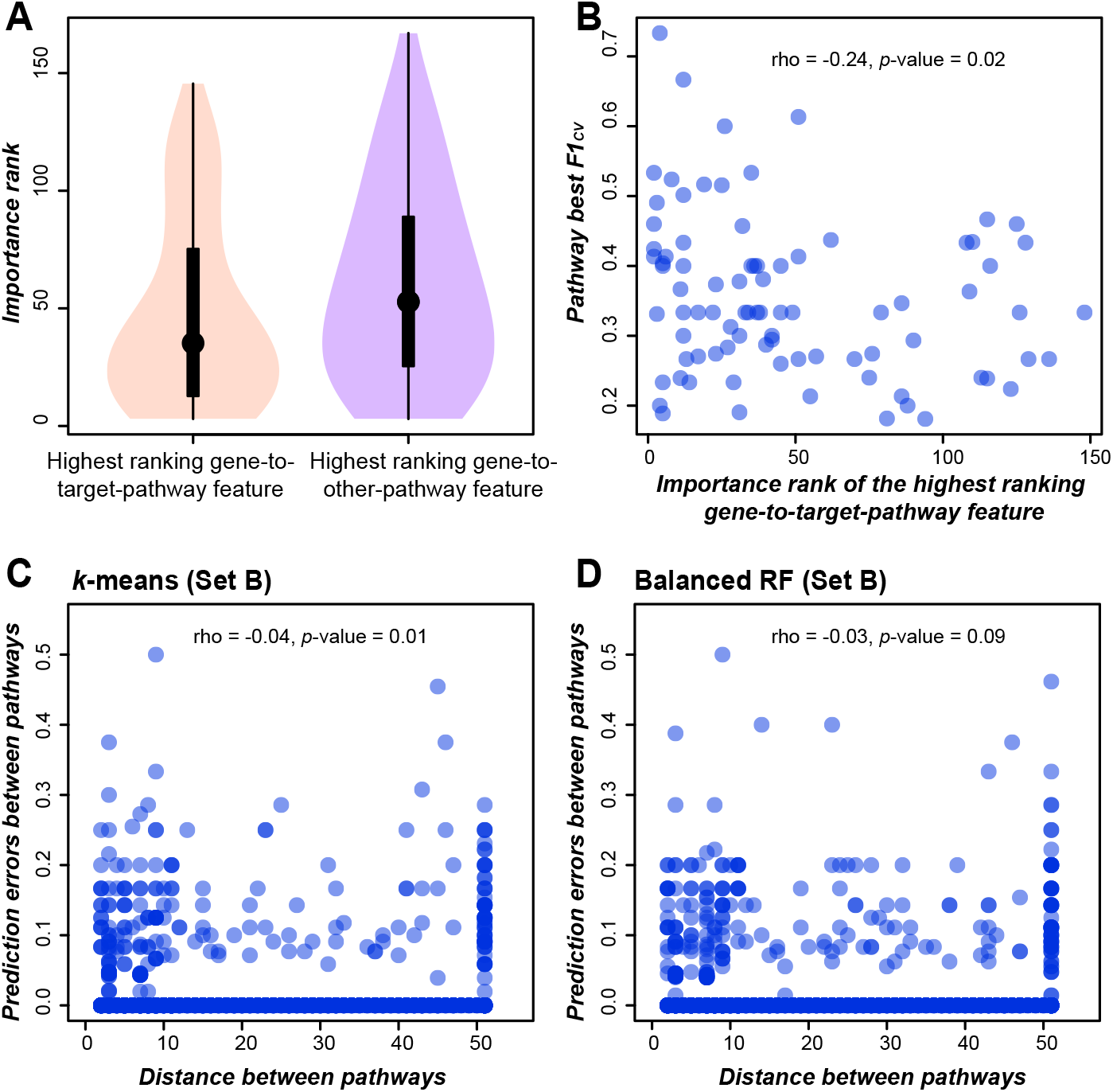
Potential reasons for prediction errors. (**A,B**) Correlation between prediction errors (proportion of genes in pathway A that were mis-predicted as being in pathway B) and the metabolic network distance between pathway A and B for *k*-means Set B models (**A**) and balanced RF Set B models (**B**). A pair of pathways have a distance of 50 if they are connected by ≥ 50 connections or have no connections. (**C**) The importance ranks of the highest ranking gene-to-target-pathway (pink) and gene-to-other-pathway (purple) features in pathway best RF Set B models. (**D**) Correlation between the importance ranks of the highest ranking gene-to-target-pathway features and the pathway best F1_CV_ in pathway best RF Set B models.

We assessed the cause of mis-predictions in two ways. First, we asked whether some genes were predicted to be in neighboring pathways based on the hypothesis that the more closely related two pathways are, the more likely that genes from these two pathways are co-regulated and, thus, harder to distinguish using gene expression data. To test this, we first connected pairs of pathways if a pair of pathways shared common reactions and constructed a pathway network. Then we determined the distance between any two pathways in the network as the minimum number of pathway nodes separating the pathways in question. We found that the prediction errors (the proportion of genes in pathway X predicted to be in pathway Y) did not correlate with distances between pathways, neither for *k*-means models (Spearman’s rho = −0.04, *p*-value = 0.01) nor for RF models (Spearman’s rho = −0.03, *p*-value = 0.09, **Figure 6C,D**).

In our second approach, we asked how potential pathway mis-annotations may contribute to mis-predictions. Upon closer examination of potentially mis-predicted genes, we noticed that some pathway steps were annotated with an unusually high number of genes. For example, the TRIACYLGLYCEROL-LIPASE-RXN reaction (EC 3.1.1.3/3.1.1.34) in the LIPAS-PWY pathway had 66 annotated genes after data filtering (see **Methods**). In addition, we found that pathways with lower F1s tended to have significantly higher average numbers of annotated enzymes per reaction (rho = −0.37, *p* = 6.3e-4, **Supplemental Figure 7**) and a higher maximum number of annotated enzymes in a single reaction (rho = −0.48, *p* = 4.3e-6, **Figure 7A**). After excluding the top eight pathways (outliers indicated in **Figure 7A,B**) with the highest number of genes annotated to a single reaction, we recalculated the feature values and carried out supervised learning again. We found that the removal of these pathways led to better, the same, and worse predictions for 43, 17, and 17 pathways, respectively (**Figure 7C** and **Supplemental Data Set 18**). Overall, the median pathway best F1_CV_ scores improved from 0.33 before removing these outlier pathways to 0.4 after their removal (*p*-value of Wilcoxon signed-rank test = 0.049). These results demonstrate the impact of annotation quality on our predictions and, as pathway annotation errors are fixed, further improvement in performance is expected. In addition, our current models can be regarded as “baseline” predictions that can be further improved.

**Figure 7.**
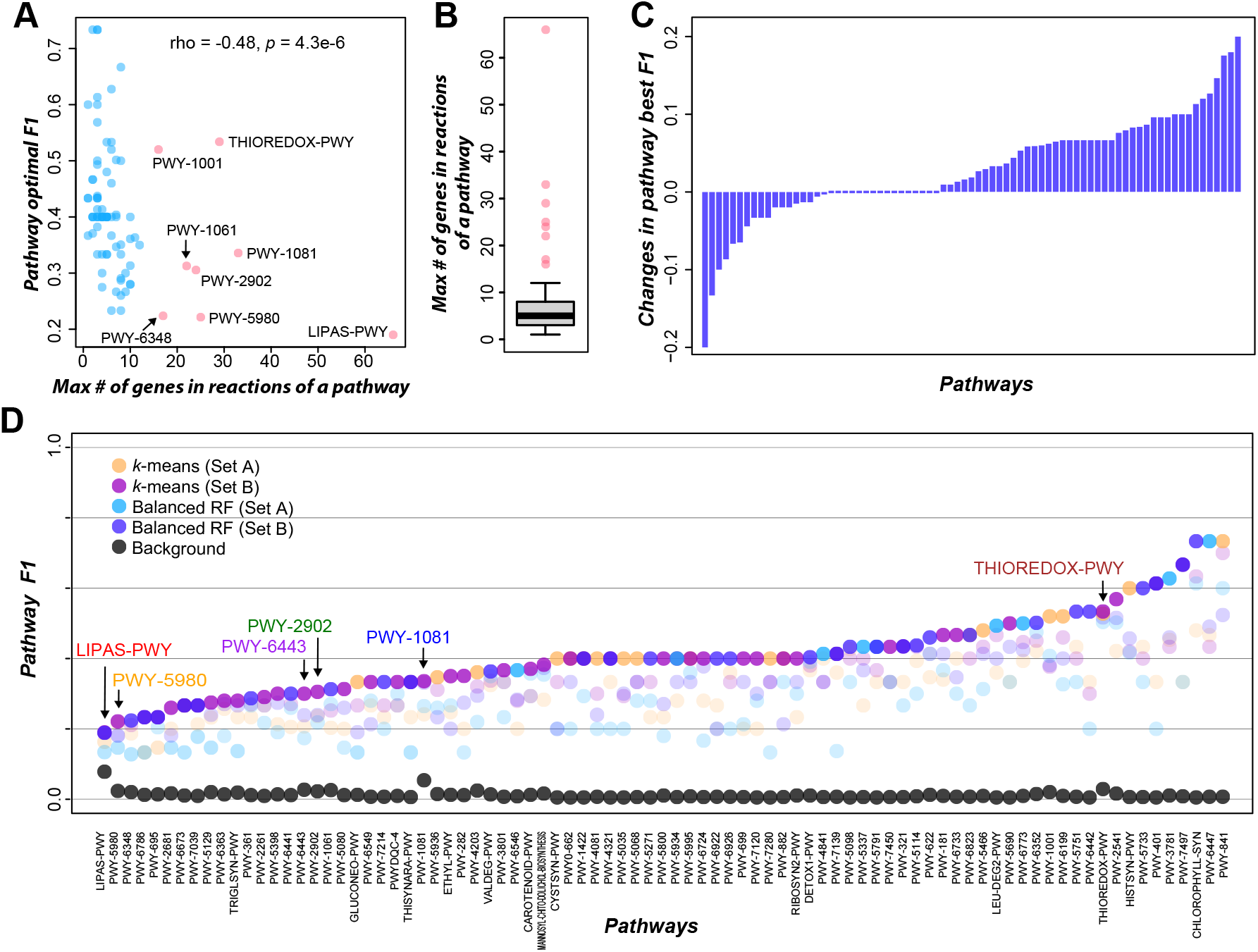
Summary of pathway membership prediction using unsupervised and supervised models. (**A**) Correlation between the maximum number of genes annotated to a single reaction in a pathway and the pathway optimal F1. Red dots: eight outlier pathways. (**B**) Boxplot of the maximum number of genes annotated to a single reaction in a pathway. Red dots: outlier pathways as in (**A**). (**C**) Prediction improvement in balanced RF Set B models when excluding eight outlier pathways. X-axis: 77 pathways, y-axis: differences in pathway F1s after excluding outlier pathways. (**D**) Pathway best F1_CV_ in *k*-means models using Set A (orange) and Set B (pink), in balanced RF models using Set A (blue) and Set B (purple), and the background pathway F1 (black). The largest six pathways with test genes are marked using different colored fonts. F1_test_ values of these six pathways in the pathway best models are shown in **Supplemental Figure 4B** and **Supplemental Data Set 20**.

### Optimizing pathway predictions by identifying optimal data and method combinations

In the previous sections, we showed the influence of prediction methods (naive prediction, *k*-means, and RF), expression measure (expression values and expression correlation), and the choice of data combination in pathway membership prediction. Because the naive prediction models performed much worse (**Figure 3C and Figure 4C**), they are not discussed here. To provide optimal predictions for different pathways, we summarized the pathway best predictions when different Set A and Set B datasets as well as different methods were used to build prediction models. The optimal F1 values ranged from 0.19 to 0.73, with a median pathway optimal F1 of 0.4 (**Figure 7D**). For a pathway with 20 members, a median F1 of 0.4 means that, to recover 10 (50%) of the genes in this pathway, 30 predictions would need to be made where 10 (33%) predictions are true positives while 20 (67%) are false positives. An F1 of 0.4 may seem low at first glance, but it is much higher compared with the median background F1 of 0.01 (the F1 measure one would achieve by randomly guessing, **Supplemental Data Set 13**). In the random guessing scenario, to recover 50% of genes in a pathway with 20 members, 2000 predictions would need to be made where only 0.5% of the predictions would be true positives while 99.5% would be false positives. Thus, while there is still considerable room for improvement, the optimal predictions are substantially better than the background.

We next asked whether optimal prediction tends to be achieved when particular data combinations are used. We found that the “condition-independent” dataset (including all experiments) provided optimal prediction for seven pathways (**Supplemental Figure 8**). The pathways that were best predicted with all experiments are those involved in general cellular processes. For example, the thioredoxin pathway (THIOREDOX-PWY) is important for maintaining intracellular redox status and essential for a plethora of downstream processes. Other examples include the TCA cycle II pathway (PWY-5690), two Fatty Acid and Lipid Biosynthesis pathways (linoleate biosynthesis I [PWY-5995], very long chain fatty acid biosynthesis I [PWY-5080]) and two cell wall biosynthesis pathways (cellulose biosynthesis [PWY-1001], xylan biosynthesis [PWY-5800]) (**Supplemental Data Set 19**).

In contrast, the remaining 78 pathways (91.8%) were better predicted using “condition-dependent” datasets (**Supplemental Figure 8**), echoing our findings on the impact of dataset on expression correlation. We also found that pathway membership tended to be better predicted when datasets for potentially related biological processes were used. For example, the abscisic acid (ABA) biosynthesis pathway (PWY-695), and the superpathway of carotenoid biosynthesis in plants (CAROTENOID-PWY), which produces carotenoids serving as precursors for ABA biosynthesis (Milborrow, 2001), had the optimal F1 when the hormone treatment dataset was used (**Supplemental Data Set 19**). Another example is the galactolipid biosynthesis I pathway (PWY-401), which had the highest F1s when samples with ABA treatment were used (**Supplemental Data Set 19**). In *Arabidopsis thaliana*, genes in PWY-401, *MONOGALACTOSYL DIACYLGLYCEROL SYNTHASE* (*MGD*) 1/2/3 and *DIGALACTOSYL DIACYLGLYCEROL DEFICIENT* (*DGD*) 1/2, were previously found to be responsive to phosphate starvation in an ABA-dependent manner (Woo et al., 2012). In addition, an optimal F1 for the trichome monoterpenes biosynthesis pathway (PWY-6447) was obtained when a pathogen treatment dataset was used (**Supplemental Data Set 19**), which supports a role of monoterpenes in the defense against pathogens (Lackus et al., 2018). Finally, the F1 for the gluconeogenesis I pathway (GLUCONEO-PWY) was optimal when the fruit ripening dataset was used (**Supplemental Data Set 19**), consistent with findings tying this pathway to fruit ripening (Famiani et al., 2016) and the observation that enzymes from this pathway exhibit differential protein abundance during fruit ripening (Pontiggia et al., 2019). These results not only illustrate the importance of the data set used, but also highlight the possibility of using the prediction framework here to identify connections between pathways and biological processes.

## Conclusion

Plant metabolites are highly diverse and play important roles in plant survival, human nutrition and medicine. Most enzyme encoding genes of plants remain unassigned to metabolic pathways. As transcriptome data have proliferated, gene coexpression analysis has been widely used to associate genes with specific functions. In this study, we aimed to assess the utility of gene expression data for predicting metabolic pathway membership by considering 82 expression values (41 expression datasets x 2 expression values) and 656 gene-to-pathway expression similarity (41 expression datasets x 2 expression values x 8 similarity measures) data combinations, and three prediction strategies (naive prediction, unsupervised and supervised learning). We demonstrated that the optimal data combination and prediction strategy should be identified for each pathway. Among the 85 pathways examined, 90 different data combinations (⍰ 1 data combinations may lead to an optimal prediction for a pathway) led to optimal membership predictions for these pathways (i.e., pathway best predictions). In addition, 59 and 39 pathway best predictions were made by unsupervised and supervised learning approaches, respectively. Notably, both machine learning approaches made significantly better pathway membership assignments compared with naive methods. Finally, our examples demonstrate that optimal pathway membership predictions tend to be achieved when pathway function-associated datasets are used. However, in most cases, it is not readily obvious why a data combination is optimal for a specific pathway. The unsupervised learning approach outperformed supervised learning when the pathway size was small, likely because there was insufficient data for supervised learning to be effective. Nonetheless, the correlation between pathway size and performance difference between these two approaches is far from perfect. These findings further indicate the importance of exploring both data combination and approach when making pathway membership predictions. As a future direction, it will be helpful to dissect the prediction models using model interpretation methods (Azodi et al., 2020) to further identify which data features are particularly important for predicting each member of a pathway; this will allow us to pinpoint the nature of useful data and develop a strategy for identifying optimal data combinations.

We also demonstrated that while the prediction performance measures (F1) of some pathways (e.g., phosphate acquisition [PWY-6348], detoxification of reactive carbonyls in chloroplasts [PWY-6786], **Figure 6A**) were much better than random guessing, the predictions had relatively high error rates no matter which data or algorithms were examined. There are three potential reasons for this. First, a good prediction model relies on the quality of the input data. In tomato, the current pathway annotation mainly relies on the sequence similarity of tomato genes to those in other model species such as *A. thaliana*, where the pathway annotations of most genes (68%, 2364 of 3475, AraCyc v 17.1 in PMN 14.0) are inferred computationally. Another consideration is that the composition of metabolites varies across species due to repeated metabolic innovation via gene duplication and subsequent sub- or neofunctionalization (Pichersky and Gang, 2000), recruitment of genes to new pathways (Shoji and Hashimoto, 2011), and loss of pathway genes due to lack of selection or selection against them in new environments (Cutter and Jovelin, 2015; Baggs et al., 2020). Thus, membership of genes in lineage-specific pathways may not be readily inferred using information from model organisms. Nonetheless, as annotation improves, the accuracy of predictions is also expected to increase.

Second, genes with multiple pathway annotations were removed in our study to simplify pathway assignment. This approach further reduced the size of the training data sets. To overcome this issue, multilabel supervised learning approaches (Herrera et al., 2016), where multiple pathways can be assigned to a gene, can be used. Third, additional features (i.e., predictive variables) may be needed to improve the pathway predictions. For example, some pathways are composed of several interwoven reactions, and genes with opposite or competitive functions may have inversely related expression profiles (Zeng and Li, 2010). In this case, the biochemical reactions that enzymes catalyze could provide more information about the interaction network within a pathway and such information could be incorporated as features. In addition, only transcriptome-based features were used in this study. Considering that enzymes in the same pathway may be located in tandem clusters (Field et al., 2011) and interact genetically and/or physically (Weissenborn and Walther, 2017), clustering and interaction data may be informative features, although in the latter case there is currently a paucity of such data available.

Taken together, our study provides quantitative measures of the usefulness of expression data in predicting metabolic pathway membership. In addition, because quantitative measures of prediction performance are provided, our findings lay the foundation for further method comparison studies that seek to improve the use of expression data for similar purposes. Although the prediction exercise here focused on using annotated enzyme genes, the *k*-means clustering and RF models can be further applied to unknown genes and provide pathway membership predictions with estimated likelihood scores. Most importantly, our study underscores the feasibility and limitations of using only gene expression data for predicting membership in metabolic pathways. In addition, the exploration of methods and data subsets in this study provides a baseline for future modeling efforts and highlights the need for further exploration, particularly of the causes of mis-predictions, for improving future predictive models.

## Methods & Materials

### Gene functional annotation

Metabolic pathway annotations of genes in tomato (*Solanum lycopersicum*) were downloaded from TomatoCyc V3.0 (Plant Metabolic Network, PMN 12.0, https://www.plantcyc.org/) (Schlapfer et al., 2017); the gene annotations were derived from *S. lycopersicum* V3.2 (Solanaceae Genomics Network, SGN, https://solgenomics.net/, referred as SGN_V3.2). To make the gene IDs consistent with those in our previous paper (Wang et al., 2018), which were taken from the *S. lycopersicum* genome in the National Center for Biotechnology Information (NCBI V2.5, referred as NCBI_V2.5), we synchronized gene names between these two versions (**Supplementary Supplemental Data Set 2**). The coding sequence of a gene in SGN_V3.2 was considered a match to an NCBI_V2.5 entry if their alignment identity score was 100% with no gaps using the BLAST-like alignment tool (BLAT) (Kent, 2002).

The functional annotation of genes was conducted using PMN Ensemble Enzyme Prediction Pipeline V3.0 (https://gitlab.com/rhee-lab/E2P2/) with default settings. In total, there are 11,036 genes with annotated Enzyme Commission (EC) numbers and/or metabolic reactions, out of which 2,395 genes are annotated to 485 metabolic pathways in TomatoCyc V3.0 (**Supplementary Supplemental Data Set 2**). Genes with no expression (FPKM=0) in all 372 experiments were excluded from further analysis, and pathways with <5 annotated genes were also excluded, resulting in 2,171 genes annotated in 297 pathways. To facilitate the pathway membership prediction, genes annotated to more than one pathway (including 1050 genes) were removed from further analysis. Pathways with < 5 genes after this filtering step were also removed, resulting in 972 genes in 85 pathways.

### RNA-seq data and processing

*S. lycopersicum* RNA-seq data from 47 studies (**Supplementary Supplemental Data Set 3 and S4**) were mapped to the genome sequences NCBI_V2.5, and calculation of FPKM and FC were performed as previously described (Wang et al., 2018). Median FPKM value among replicates was used as the expression level for a gene in an experiment. The experiments/samples from the 47 RNA-seq studies were classified into four categories: 1) tissues/stages/circadian—samples from wild-type tomato plants taken from different tissues, at different development stages, or at different times of day; 2) genetic background—samples from comparison studies of wild type, gene mutant, or transgenic plants where expression of a gene was knocked down or overexpressed; 3) hormone treatment—samples from plants treated with hormones (cytokinin, ABA, auxin, etc.) or control treatments; 4) stress treatment—samples from plants with stress (pathogen, light, cold, heat, etc.) or control treatments (**Supplementary Supplemental Data Set 3**). Samples belonging to each of these four categories constituted a collection of expression experiments referred to as a compiled dataset, thus resulting in four compiled datasets. A fifth compiled dataset was made by combining all 372 experiments. In addition, only 36 of these 47 individual datasets had ⍰ 3 experiments and were used further as independent individual datasets. Thus, there were 41 datasets in total: 36 individual datasets and five compiled datasets.

### Gene expression similarity measure

Eight similarity measures were evaluated in our study. The first two, Pearson correlation coefficient (PCC) and Spearman’s rank correlation coefficient (Spearman) between expression values of two genes, were calculated using the pearsonr and the spearmanr functions, respectively, in the scipy.stats module (Virtanen et al., 2020). The third measure, Mutual Information (MI) was determined using the normalized_mutual_info_score function in the python module sklearn.metrics.cluster (Pedregosa et al., 2011). Partial correlation (partCor), the fourth measure, was determined with the pcor.shrink function (Schafer and Strimmer, 2005) in the corpcor R package. In addition, Mutual Rank (MR) was calculated for each of these four measures. For a pair of genes, *gene1* and *gene2*, the MR based on a particular similarity measure *S* is

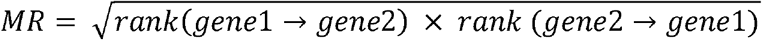

Where *rank_(gene1->gene2)_* and *rank_(gene2->gene1)_* indicate the rank of expression similarity between *gene1* and *gene2* among all expression similarity values calculated between *gene2* and other genes and between *gene1* and other genes, respectively. The ranks range from 1 to 2,170, with 1 indicating highest expression similarity. Thus, including all four MR-based measures, there were eight expression similarity measures examined.

### Data combinations

Two types of data combinations were used in the pathway membership predictions. The first type (Set A) contained gene expression values in an experiment (FPKM) or fold changes between experiments (FC). Thus, there were 82 Set A data combinations (41 expression datasets X 2 expression values). For each Set A data combination, the number of features equaled the number of experiments (FPKM) or the number of experiment comparisons (FC) in the expression dataset. The second type (Set B) contained median and maximum expression similarities between a gene and genes in a pathway. The gene-to-pathway expression similarities were calculated using eight similarity measures, thus resulting in 656 Set B data combinations (41 X 2 X 8). For each Set B data combination, there were 170 features (2 similarity values X 85 pathways).

### Splitting data for testing and modeling, and measuring prediction performance

To assess how well the approaches predicted pathway membership, five genes from each of the six pathways with ≥ 25 genes expressed in ≥ 1 samples were held out as test data for evaluating unsupervised and supervised approaches. Note that these hold-out genes were not used in any cluster/model building process. The remaining genes in these six pathways as well as all genes from the remaining 79 pathways were split into five groups. Genes in four of the five groups were referred to as “training” genes for all three approaches: naive prediction, unsupervised, and supervised. Genes in the remaining, fifth, group were used as “validation” genes to evaluate the prediction performance of those three approaches. This split was repeated five times to ensure that every gene was placed in a validation subset once.

F-measure (F1) was calculated by comparing the annotated and predicted pathway membership for validation genes in a pathway (referred as F1_CV_), using the equation: 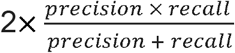, where 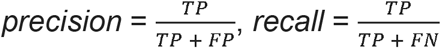: true positive, i.e., number of genes annotated and predicted as being a member of the pathway. *FP*: false positive, i.e., number of genes mis-predicted as being in the pathway. *FN*: false negative, i.e., number of genes annotated but not predicted as being in the pathway. Average F1_CV_ for each pathway across the five data splits mentioned above were used to evaluate prediction performance.

### Naive prediction

Two different pathway membership predictions were made using the naive prediction strategy. First, gene X was predicted to be in pathway A if gene X had the highest median expression similarity with genes in pathway A compared with those in other, non-A pathways. This method was referred to as naive median prediction. Second, gene X was predicted to be in pathway B if gene X had the maximum expression similarity with gene Y, which is in pathway B. This method was referred to as naive maximum prediction. A gene was not predicted to be in any pathway if it had ≥ 2 pathway assignments in naive median or naive maximum prediction.

### Unsupervised learning

The unsupervised clustering was conducted using the “*k*-Means” and “AffinityPropagation” functions in sklearn.cluster (Pedregosa et al., 2011). For *k*-means, the parameter *n_clusters* (how many clusters to get, i.e., *k*) at 5, 10, 25, 50, 85, 100, 200, 300, 400 and 500 was tested. For Affinity Propagation, damping factor, i.e., the extent to which the current value is maintained relative to incoming values, at 0.5, 0.6, 0.7, 0.8, 0.9 and 0.99 were used. To evaluate if genes in a cluster X were enriched among pathway A genes, an enrichment value for A and X was defined as: 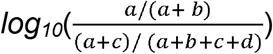: (*a*) the numbers of genes in A and X, (*b*) the number of genes in A but not in X, (*c*) the number of genes in X but not in A, (*d*) the number of genes in neither A nor X. Genes in cluster X were assigned to pathway A if the enrichment value for A and X was the highest among enrichment values for X and other pathways, and the *p*-value of corresponding Fisher’s exact test with *a*, *b*, *c*, and *d* in a 2×2 contingency table was <0.05. For validation genes not used for generating clusters, the cluster memberships were predicted using the “predict” function in sklearn.cluster. Since there were 82 Set A and 656 Set B data combinations, the above process was repeated for each data combination to identify the dataset that led to clusters that provided the best predictions (i.e., highest F1_CV_) for each pathway. The same prediction procedure was used to predict pathway memberships for test genes.

### Supervised learning

Multi-class Random Forest (RF) models were established using training genes, with the sklearn.ensemble.RandomForestClassifier (Pedregosa et al., 2011). Hyperparameters (*max_depth [3, 5, 10], max_features [0.1, 0.25, 0.5, 0.75, ‘sqrt’, ‘log2’, None]*, and *n_estimators [100,500,1000]*) were determined by performing a grid search using the five cross-validation scheme, which was explained in the data split section, with the goal of maximizing F1_CV_ for each of the 82 Set A or 656 Set B data combinations. A final model for the data combination in question was built with the best hyperparameter identified and applied to the validation genes. Because a five-fold cross-validation scheme was used, the reported F1_CV_ for the final model for a specific data combination was the average of F1s from all five folds. To balance the numbers of genes among pathways, the training data of smaller pathways (i.e., those < 56) were up-sampled to 56 genes per pathway using the SMOTE function from imblearn.over_sampling (Blagus and Lusa, 2013), with sampling_strategy=‘not majority’, random_state=42, k_neighbors=3. The “predict” function from sklearn.ensemble.RandomForestClassifier was used to predict the pathway membership of test genes. In the same way as for the clustering-based approach, a final model was built with each of 82 Set A or 656 Set B data combinations to identify the data combination that led to the optimal prediction model for each pathway. The impurity-based feature importance was used to compare the relative degrees of contribution of features to the RF model—the higher the feature importance, the higher the relative contribution to the models—and was determined using the attribute feature_importances_ of RandomForestClassifier with criterion=gini.

## Supplementary figure legends

**Supplemental Figure 1.** Differences in gene expression similarity within pathways and between pathways.

(**A**) Maximum PCC between a gene and all other genes within the same pathway; the expression levels (FPKM) of genes in all 372 samples were used to calculate the PCC. Blue line: median value; light blue box: interquartile range. Violin plot shows distribution of maximum PCC between a gene and genes from a different pathway. Median value and interquartile range in the between-pathway distribution are marked with red and orange dashed lines, respectively.

(**B**) Scatter plot of the percentile_BP_ of median (x-axis) vs. maximum (y-axis) PCC between a gene and other genes within the same pathway.

(**C**) Distribution of PCC values between gene pairs within a pathway and between pathways. X-axis: PCC values for different within-pathway percentiles. All *p*-values are from Wilcoxon rank sum test < 5.3e-10. es: effect size, which is the difference between medians.

(**D**) Distribution of PCC values between homologous (blue) and non-homologous (light blue) gene pairs within the same pathway. X-axis: different within-pathway percentiles. Pink dashed line: median between-pathway PCC values for gene pairs in (**C**).

**Supplemental Figure 2.** Impact of similarity measure on expression similarity.

(**A**) Three examples showing pathways with different expression similarity percentile_BP_ calculated with different similarity measures. Color scale: five ranges of expression similarity percentile_BP_.

(**B**) Expression profiles (normalized FPKM) in all 372 samples for the three pathways shown in (**A**). The expression levels are scaled to 0–1, minimum level: 0 (blue), maximum: 1 (red).

**Supplemental Figure 3.** Confusion matrix for naive prediction models.

Confusion matrix showing the proportion of genes that are predicted in each pathway for pathway best naive median (**A**) and pathway best naive maximum (**B**) prediction models. Color scale: proportion of genes in a pathway (y-axis) predicted as being in one of the 85 pathways (x-axis) by the pathway best model.

**Supplemental Figure 4.** Pathway membership prediction using unsupervised and supervised models.

(**A**) Distribution of average F1_CV_ values for unsupervised (*k*-means and Affinity Propagation) and supervised models (Random Forest).

(**B**) F1_CV_ and F1_test_ (F1 value for the five test genes) values for six pathways, from which five genes were held out as test genes; values are shown for models that yielded the highest F1_CV_ for that pathway. Dots: individual pathways.

(**C**) Pathway F1_CV_ for the unsupervised and RF models that had the highest average F1CV (x-axis) and the pathway best F1CV (y-axis).

**Supplemental Figure 5.** Potential underlying reasons for the relatively poorer performances of Set A and RF models than Set B and *k*-means models, respectively. (**A,B**) Correlation between the numbers of features in Set A and pathway best F1_CV_ values from *k*-means (**A**) or RF models (**B**) when Set A was used.

(**C,D**) Correlation between the number of genes in a pathway and difference between pathway best F1_CV_ for RF and *k*-means models when Set A (**C**) or Set B (**D**) was used. (**E,F**) Same as (**C,D**), except that two outlier data points were removed.

**Supplemental Figure 6.** Pathway F1_CV_ values from *k*-means models and balanced RF models.

(**A**) Heatmap showing normalized pathway F1_CV_ for 656 *k*-means Set B models.

(**B**) Heatmap showing normalized pathway F1_CV_ for 656 RF models. The maximum F1_CV_ of a pathway was set as 1 and the minimal F1CV as 0.

**Supplemental Figure 7.** Potential reasons for low pathway F1.

(**A**) Correlation between the pathway optimal F1_CV_ and the number of annotated enzymes per reaction in a pathway. Red dots: outlier pathways. Red font: correlation after outlier pathways were removed.

(**B**) Confusion matrix of balanced RF Set B models when eight pathways were excluded. Color scale: proportion of genes in a pathway (y-axis) predicted as being in one of the 77 pathways (x-axis) in pathway best RF models.

**Supplemental Figure 8.** Pathway optimal expression datasets.

Distribution of pathway optimal F1_CV_ (y-axis, highest pathway F1_CV_ from all four methods in **Figure 7D**) when different datasets were used (x-axis). Blue dots: individual pathways.

## Funding

This work was partly supported by the National Science Foundation IOS-1546617 to C.S.B. and S.-H.S and NSF DEB-1655386 and U.S. Department of Energy Great Lakes Bioenergy Research Center BER DE-SC0018409 to S.-H.S. C.S.B is supported in part by Michigan AgBioResearch and through USDA National Institute of Food and Agriculture Hatch project number MICL02552.

## Acknowledgments

We thank John P. Lloyd, Christina B. Azodi, Serena G. Lotreck, Elizabeth M. Gibbons, Koichi Sugimoto, Yann-Ru Lou, Bryan Leong, Brian St. Aubin, Pengxiang Fan, Paul Fiesel, Craig Schenck, Matthew Bedewitz, Robert Last, and Eran Pichersky for helpful discussions.

## Computational resources

All the scripts used in this study are available on Github at: https://github.com/ShiuLab/Pathway_gene_prediction_in_tomato

## Data availability

Set A and Set B data are available on Zenodo at: https://zenodo.org/record/4050240#.X3TufDNvrrN

## Author contributions

P.W. and S.-H.S conceived and designed the study. P.W., B.M.M., and S.U. performed the analysis. P.W., M.D.L., C.B. and S.-H.S. wrote the paper. All authors read and approved the final manuscript.

## Notes

### Competing Interest Statement

The authors have declared no competing interest.

