## Supplemental figures for "Optimizing the use of gene expression data to predict plant metabolic pathway memberships"

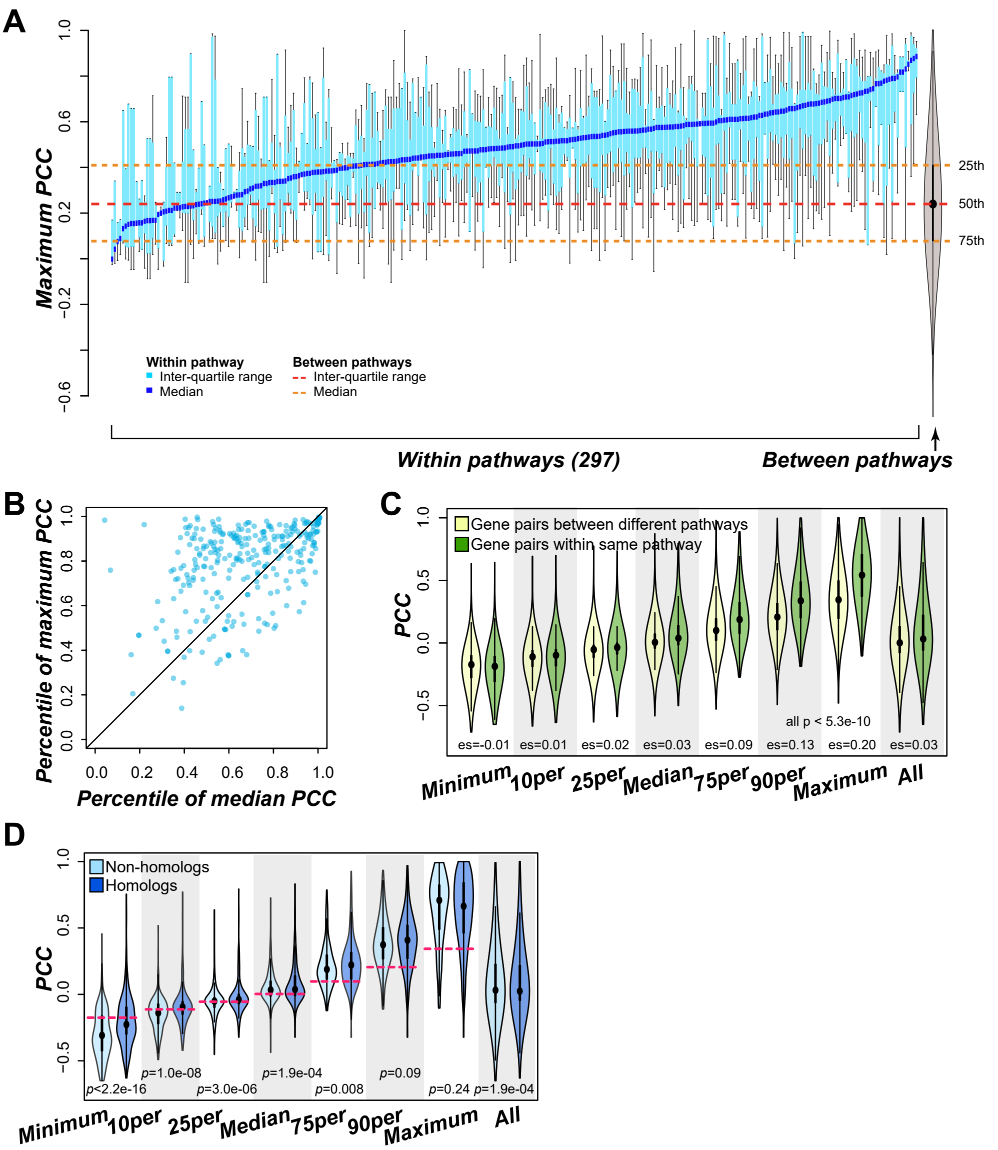


**Supplemental Figure 1.** Differences in gene expression similarity within pathways and between pathways.

(**A**) Maximum PCC between a gene and all other genes within the same pathway; the expression levels (FPKM) of genes in all 372 samples were used to calculate the PCC. Blue line: median value; light blue box: interquartile range. Violin plot shows distribution of maximum PCC between a gene and genes from a different pathway. Median value and interquartile range in the between-pathway distribution are marked with red and orange dashed lines, respectively.
(**B**) Scatter plot of the percentileBP of median (x-axis) vs. maximum (y-axis) PCC between a gene and other genes within the same pathway.
(**C**) Distribution of PCC values between gene pairs within a pathway and between pathways. X-axis: PCC values for different within-pathway percentiles. All *p*-values are from Wilcoxon rank sum test < 5.3e-10. es: effect size, which is the difference between medians.
(**D**) Distribution of PCC values between homologous (blue) and non-homologous (light blue) gene pairs within the same pathway. X-axis: different within-pathway percentiles. Pink dashed line: median between-pathway PCC values for gene pairs in (**C**).

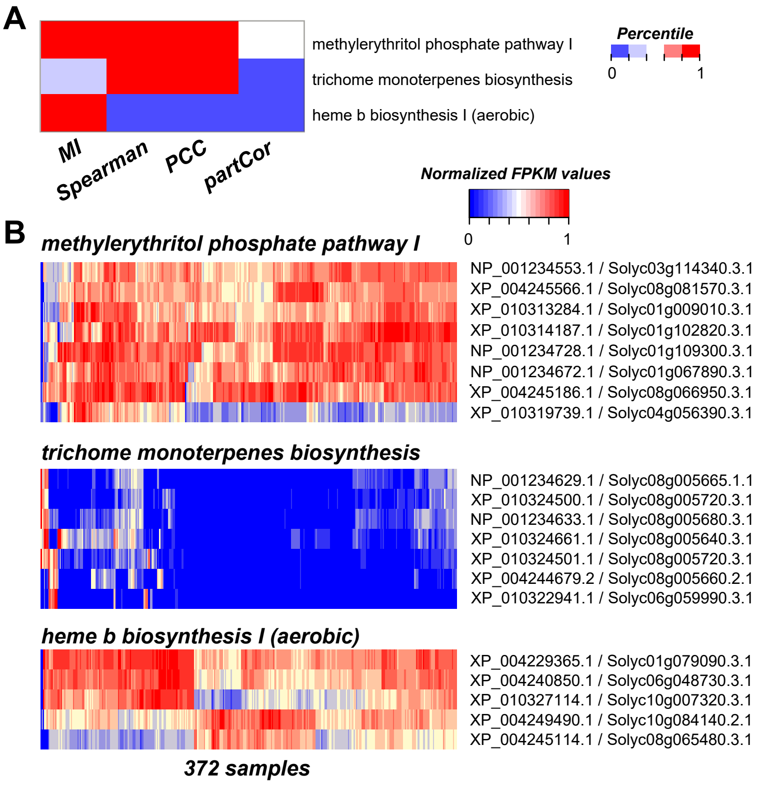


**Supplemental Figure 2.** Impact of similarity measure on expression similarity.

(**A**) Three examples showing pathways with different expression similarity percentile$_{BP}$ calculated with different similarity measures. Color scale: five ranges of expression similarity percentile$_{BP}$.
(**B**) Expression profiles (normalized FPKM) in all 372 samples for the three pathways shown in (**A**). The expression levels are scaled to 0–1, minimum level: 0 (blue), maximum: 1 (red).

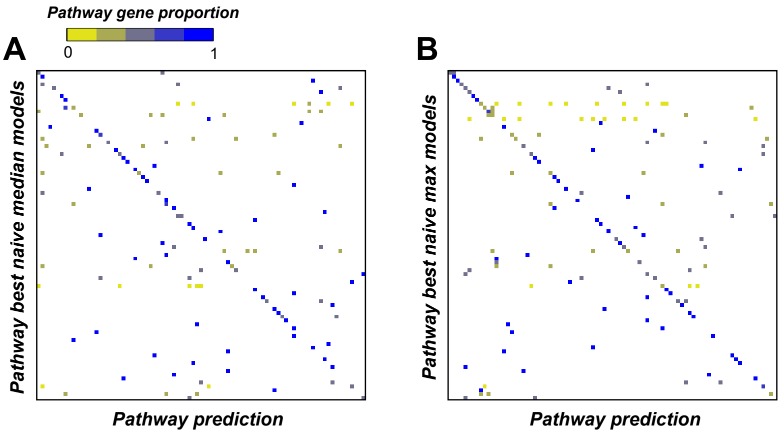


**Supplemental Figure 3.** Confusion matrix for naive prediction models.

Confusion matrix showing the proportion of genes that are predicted in each pathway for pathway best naive median (**A**) and pathway best naive maximum (**B**) prediction models. Color scale: proportion of genes in a pathway (y-axis) predicted as being in one of the 85 pathways (x-axis) by the pathway best model.

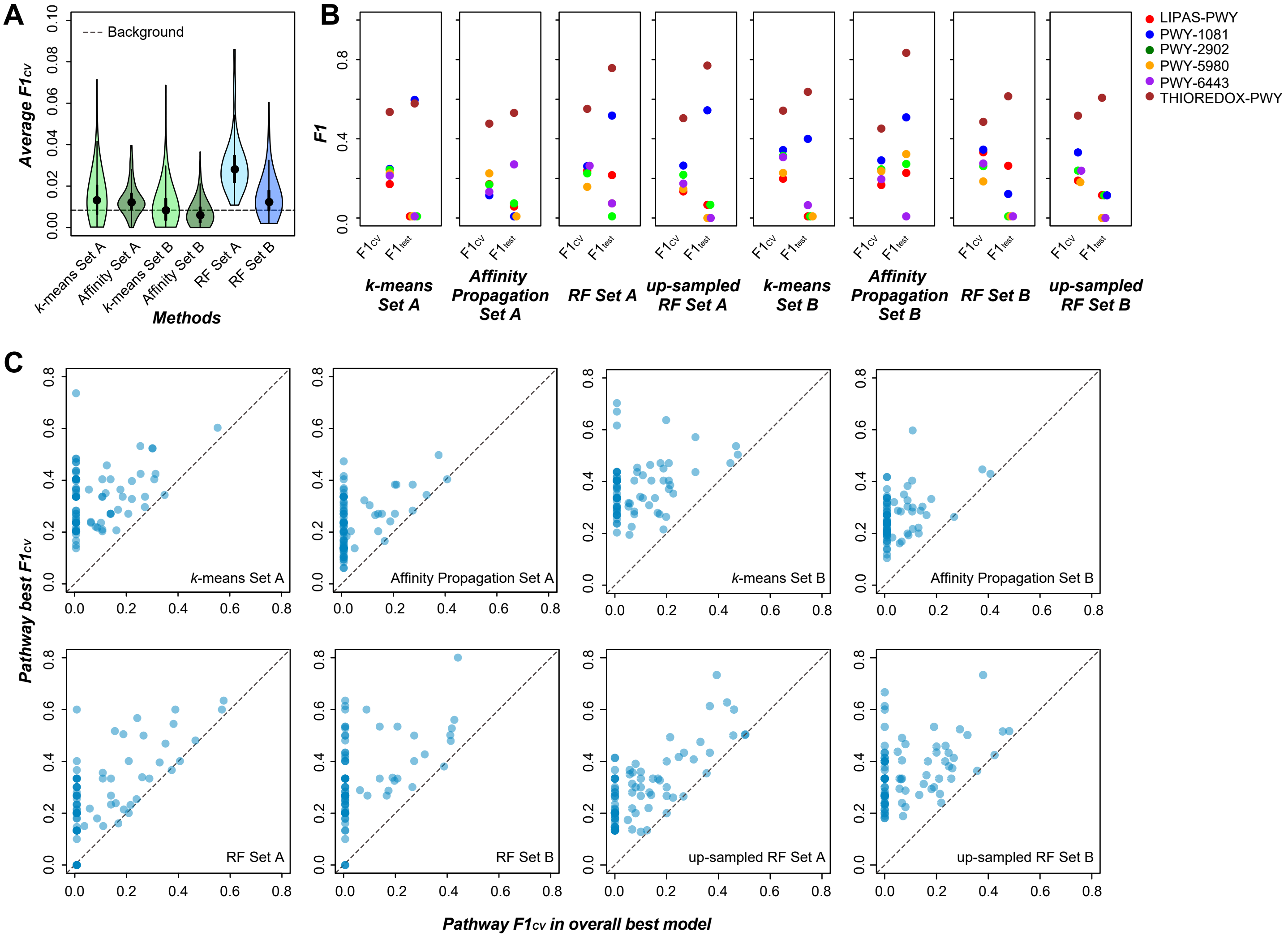


**Supplemental Figure 4.** Pathway membership prediction using unsupervised and supervised models.

(**A**) Distribution of average $F1_{cv}$ values for unsupervised (*k*-means and Affinity Propagation) and supervised models (Random Forest).
(**B**) $F1_{cv}$ and $F1_{test}$ (F1 value for the five test genes) values for six pathways, from which five genes were held out as test genes; values are shown for models that yielded the highest $F1_{cv}$ for that pathway. Dots: individual pathways.
(**C**) Pathway $F1_{cv}$ for the unsupervised and RF models that had the highest average $F1_{cv}$ (x-axis) and the pathway best $F1_{cv}$ (y-axis).

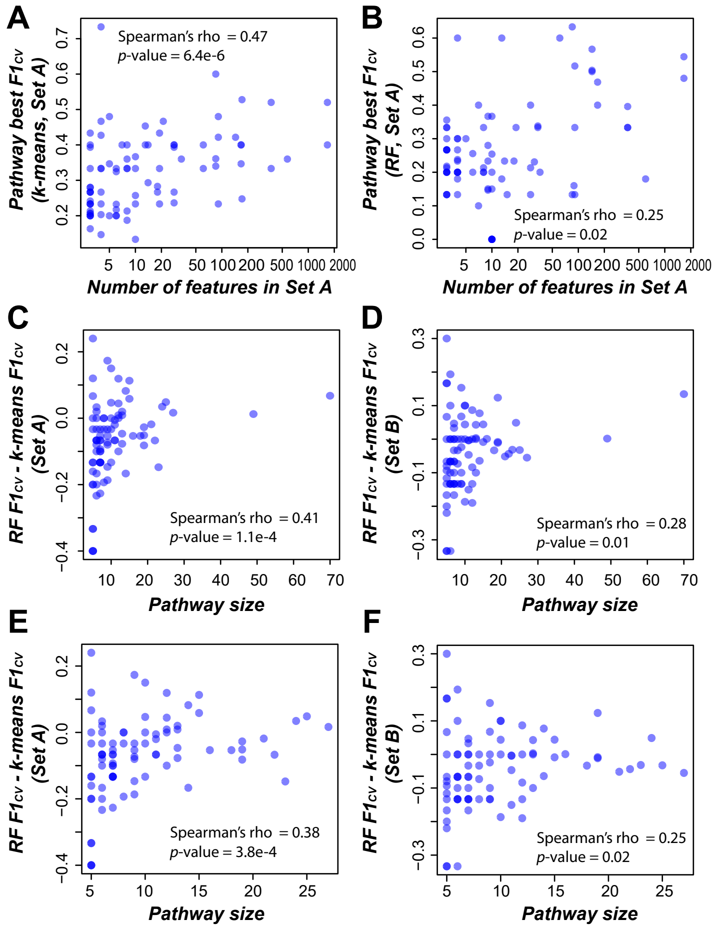


**Supplemental Figure 5.** Potential underlying reasons for the relatively poorer performances of Set A and RF models than Set B and *k*-means models, respectively.

(**A,B**) Correlation between the numbers of features in Set A and pathway best $F1_{cv}$ values from k-means (**A**) or RF models (**B**) when Set A was used.
(**C,D**) Correlation between the number of genes in a pathway and difference between pathway best $F1_{cv}$ for RF and *k*-means models when Set A (**C**) or Set B (**D**) was used.
(**E,F**) Same as (**C,D**), except that two outlier data points were removed.

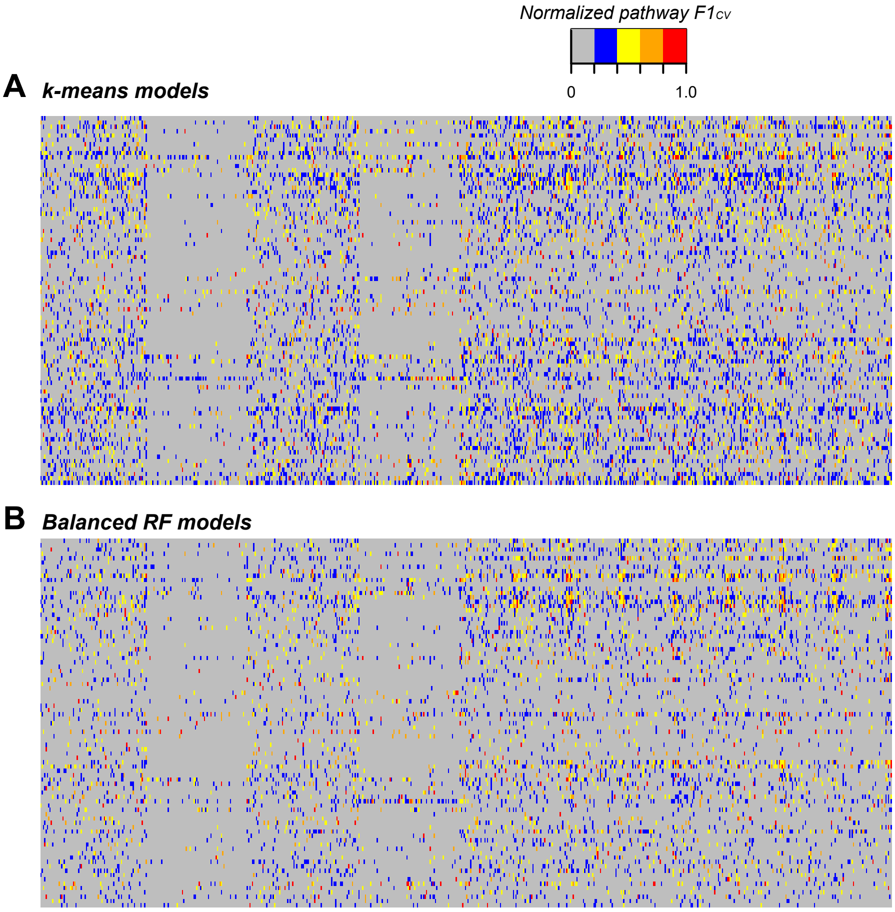


**Supplemental Figure 6.** Pathway $F1_{cv}$ values from *k*-means models and balanced RF models.

(**A**) Heatmap showing normalized pathway $F1_{cv}$ for 656 *k*-means Set B models.
(**B**) Heatmap showing normalized pathway $F1_{cv}$ for 656 RF models. The maximum $F1_{cv}$ of a pathway was set as 1 and the minimal $F1_{cv}$ as 0.

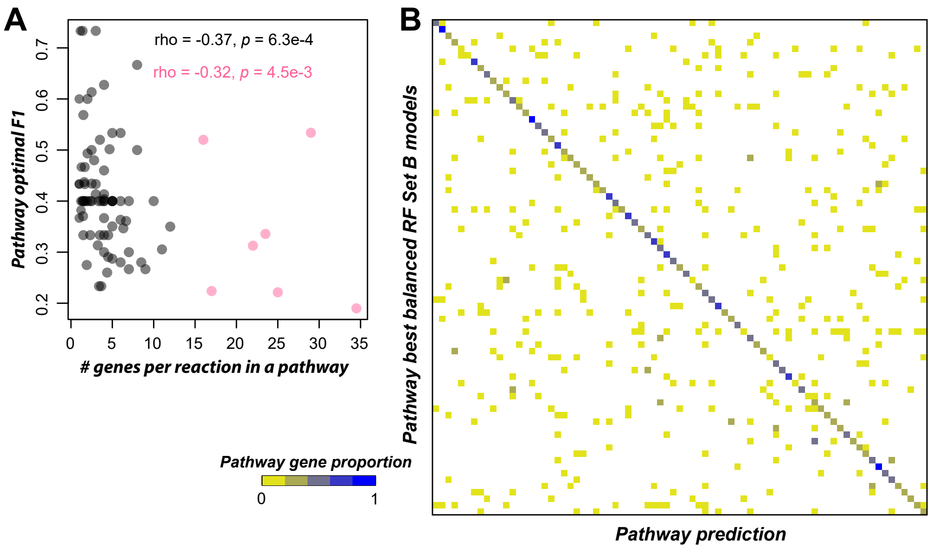


**Supplemental Figure 7.** Potential reasons for low pathway F1.

(**A**) Correlation between the pathway optimal $F1_{CV}$ and the number of annotated enzymes per reaction in a pathway. Red dots: outlier pathways. Red font: correlation after outlier pathways were removed. (**B**) Confusion matrix of balanced RF Set B models when eight pathways were excluded. Color scale: proportion of genes in a pathway (y-axis) predicted as being in one of the 77 pathways (x-axis) in pathway best RF models.

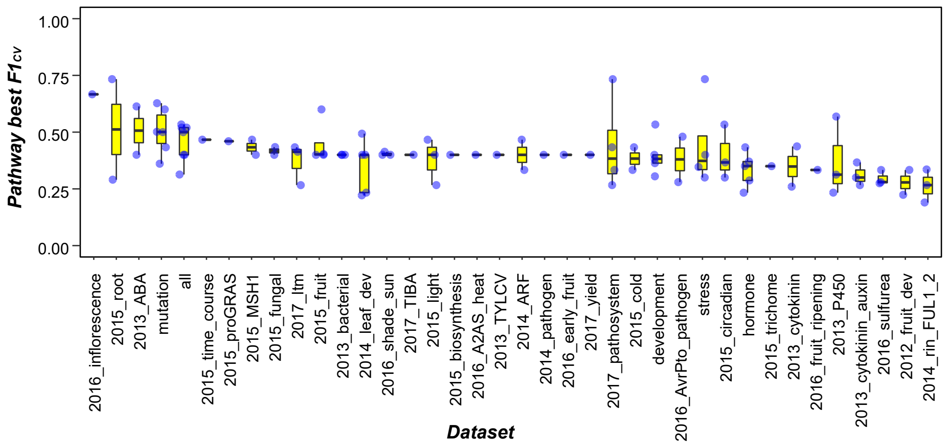


**Supplemental Figure 8.** Pathway optimal expression datasets.

Distribution of pathway optimal $F1_{CV}$ (y-axis, highest pathway $F1_{CV}$ from all four methods in **Figure 7D**) when different datasets were used (x-axis). Blue dots: individual pathways.
